# Identifying metabolic adaptations during cardiotoxicity using paired transcriptomics and metabolomics data integrated with a model of heart metabolism

**DOI:** 10.1101/2021.10.31.466544

**Authors:** Bonnie V. Dougherty, Kristopher D. Rawls, Matthew L. Jenior, Bryan Chun, Sarbajeet Nagdas, Jeffrey J. Saucerman, Glynis L. Kolling, Anders Wallqvist, Jason A. Papin

**Affiliations:** Department of Biomedical Engineering, University of Virginia, Charlottesville, VA, 22908, USA; Department of Microbiology, Immunology, and Cancer Biology, University of Virginia Health System, Charlottesville, VA 22908, USA; Department of Medicine, Division of Infectious Diseases and International Health, University of Virginia, Charlottesville, VA, 22908, USA; Department of Defense Biotechnology High Performance Computing Software Applications Institute, Telemedicine and Advanced Technology Research Center, U.S. Army Medical Research and Development Command, Fort Detrick, Maryland, 21702 USA; Department of Biochemistry & Molecular Genetics, University of Virginia, Charlottesville, VA, 22908, USA

**Keywords:** cardiotoxicity, genome-scale metabolic network reconstruction, chemotherapy-induced cardiotoxicity

## Abstract

Improvements in the diagnosis and treatment of cancer has revealed the long-term side effects of chemotherapeutics, particularly cardiotoxicity. Here, we present paired transcriptomics and metabolomics data characterizing in vitro cardiotoxicity to three compounds: 5-fluorouracil, acetaminophen, and doxorubicin. Standard gene enrichment and metabolomics approaches identify some commonly affected pathways and metabolites but are not able to readily identify metabolic adaptations in response to cardiotoxicity. The paired data was integrated with a genome-scale metabolic network reconstruction (GENRE) of the heart to identify shifted metabolic functions, unique metabolic reactions, and changes in flux in metabolic reactions in response to these compounds. Using this approach, we confirm known mechanisms of doxorubicin-induced cardiotoxicity and provide hypotheses for metabolic adaptations in cardiotoxicity for 5-fluorouracil, doxorubicin, and acetaminophen.

## Introduction

Multiple chemotherapeutics have been identified as increasing the incidence of cardiovascular events, both in the short- and long-term following treatment, now termed cardiotoxicity (Albini et al, 2010). It is now well-established that multiple chemotherapeutics are associated with adverse cardiovascular events, such as left ventricular dysfunction and chronic heart failure (Albini et al, 2010). However, mechanisms of cardiotoxicity are not well understood. Recently, changes in glucose uptake have been noted to precede clinical measures of heart dysfunction in both spontaneously hypertensive rats (Li et al, 2019) and in patients undergoing chemotherapy with known cardiotoxic drugs (Bauckneht et al, 2017; Borde et al; 2012), suggesting that broad metabolic adaptations to drug treatment may yield insight into mechanisms of cardiotoxicity.

Genome-scale metabolic network reconstructions (GENREs) provide an opportunity to mechanistically connect changes in metabolites with changes in transcriptomics, identifying potential mechanisms for the production of metabolites. GENREs provide a mechanistic representation of cellular metabolism, including the stoichiometric coefficients for metabolic reactions and the connectivity between genes and the individual reactions they govern. Previous studies have used transcriptomics data with metabolic network reconstructions to study metabolic adaptations in hepatotoxicity (Rawls et al, 2018; Pannala et al, 2018; Pannala et al, 2019; Pannala et al, 2020) and nephrotoxicity (Rawls et al, 2020; Pannala et al, 2020).

In the current study, we extend that work using a new approach to integrate paired transcriptomics and metabolomics data with a heart-specific genome-scale metabolic network (GENRE) (Dougherty et al, 2020) to predict metabolic adaptations in cardiotoxicity. Paired transcriptomics and metabolomics data was collected for primary rat neonatal cardiomyocytes exposed to three compounds: 5-fluorouracil (5FU), acetaminophen (Ace), and doxorubicin (Dox). Both Dox and 5FU were selected based on their established cardiotoxicity (Volkova et al, 2013; Sara et al, 2018) while Ace was chosen based on previous studies exploring hepatotoxicity and nephrotoxicity (Rawls et al, 2018; Rawls et al, 2020). Furthermore, Dox has multiple hypothesized mechanisms of toxicity (Volkova et al, 2013) whereas 5FU does not have established hypotheses for mechanisms of cardiotoxicity (Sara et al, 2018). Integrating multiple forms of omics data with functional models of metabolism yields unique insight into potential metabolic adaptations in cardiotoxicity. Through this integrated approach, we (a) recapitulate published metabolic adaptations in 5FU and Dox cardiotoxicity, (b) propose new metabolic adaptations in Ace cardiotoxicity, and (c) propose the role of shifts in key metabolic reactions as representative of increased metabolic demand in 5FU and Dox in response to the primary chemotherapeutic mechanisms of action.

## Results

### *Optimizing concentrations of compounds to characterize* in vitro *cardiotoxicity*

Given that most toxicity studies have limited rationale for their chosen drug concentrations, we aimed to deliberately choose cardiotoxic concentrations of 5-fluorouracil (5FU), acetaminophen (Ace), and doxorubicin (Dox) for our *in vitro* studies. Cardiotoxic concentrations were defined as the concentration that elicited a significant decrease in cell reducing potential without a significant increase in cell death compared to the control at 24 hours (Figure 1). 10 mM 5FU and 2.5 mM Ace elicited a significant decrease in cell reducing potential at 24 hours without a significant increase in cell death. 1.25 µM Dox induced a significant decrease in cell reducing potential but also a significant increase in cell death at 24 hours (Figure 1). To confirm a change in metabolism and not simply an increase in cell death, the Seahorse mitochondrial stress test was used, where the measured OCR was normalized to live cell counts (Supplemental Figure 1). There was a significant increase in OCR for ATP production for 10 mM 5FU and 1.25 µM Dox and a significant increase in basal respiration for 1.25 µM Dox. This confirms a metabolic stress on a per cell level, seen as an increase in the flux of oxygen used in the electron transport chain (ATP production) and an increase in the basal rate of oxygen consumption (basal respiration).

**Figure 1.**
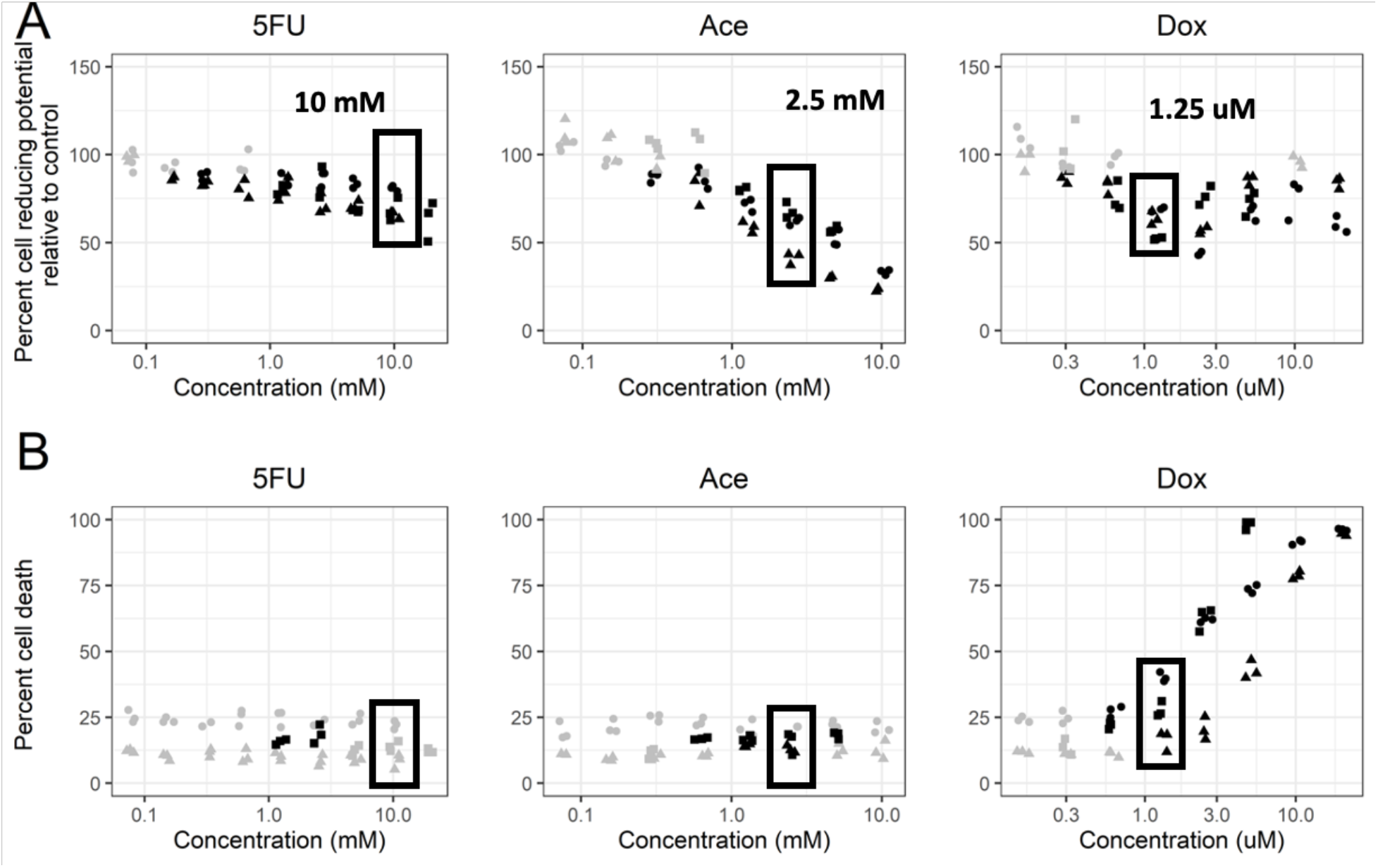
Choosing cardiotoxic drug concentrations that elicited a change in metabolism without a significant increase in cell death. (A) Percent cell reducing potential with respect to controls was measured for a range of concentrations for the chosen compounds. The shapes represent different biological replicates. Black dots indicate a statistically significant change from the control condition, determined using Dunnett’s test with a p-value < 0.05. Boxes indicate the concentrations chosen for the experimental studies. (B) Percent cell death calculated using a PI/Hoescht stain. The shapes represent different biological replicates. The black dots indicate a statistically significant change from the control condition, calculated using Dunnett’s test with p-value < 0.05.

As with previous studies (Rawls et al, 2018; Rawls et al, 2020), paired transcriptomics and metabolomics data was collected at both 6 (Supplemental Figure 2) and 24 hours (Figure 1) after drug exposure to capture early and late toxicity. There was no significant increase in cell death for any compounds for our chosen concentrations at 6 hours. There was a significant decrease in cell reducing potential for the chosen concentration of Ace at 6 hours, indicating that Ace has an early but sustained, effect on cell metabolism in cardiomyocytes (Supplemental Figure 2).

### Unsupervised machine learning of transcriptomics and metabolomics data highlights underlying drug-induced shifts in cellular activity

In order to identify the largest sources of variability between the conditions, individual samples for either transcriptomics (transcript counts) or metabolomics (metabolite abundances) data were clustered in an unsupervised fashion using Principal Component Analysis (PCA) (Figure 2). For reference, the 5FU condition is paired with the DMSO1 control and the Ace and Dox conditions are paired with the DMSO2 control (Methods). For the transcriptomics data, within each time point, the control conditions (DMSO1 and DMSO2) separate from the treatment groups (5FU, Dox, and Ace) (perMANOVA for treatment vs control, p-value < 0.001 for both 6 and 24 hour timepoints) (Supplemental Figure 3A). The first principal component separates by the Dox treatment and the second principal component separated samples by time, 6 versus 24 hours. A gene enrichment analysis of the top 100 genes in the first principal component using the Hallmark pathways from the Molecular Signatures Database (MSigDb) (Subramania et al, 2005; Liberzon et al, 2016) identified the p53 pathway as the only significantly enriched pathway, suggesting that Dox induces a unique response trhough the p53 pathway compared to the other compounds. The main function of the p53 pathway is to monitor cell division, including DNA replication (Harris et al, 2005). Two of the proposed chemotherapeutic mechanisms of action of Dox are intercalation with DNA and the inhibition of topoisomerase II (Taymaz-Nikerel et al, 2018); both mechanisms would therefore elicit a response from the p53 pathway. The second principal component separates between the 6-hour and 24-hour samples, including the control samples, suggesting a potential adaptation to culture conditions. A gene enrichment analysis for the top 100 genes in the second principal component identifies Myc targets as uniquely enriched, where consistent decreased expression of the Myc pathways is consistent with a transition from the fetal gene program (Jackson et al, 1990) and increased expression consistent with a response to cardiomyocyte stress (Lee et al, 2009; Ahuja et al, 2010).

**Figure 2.**
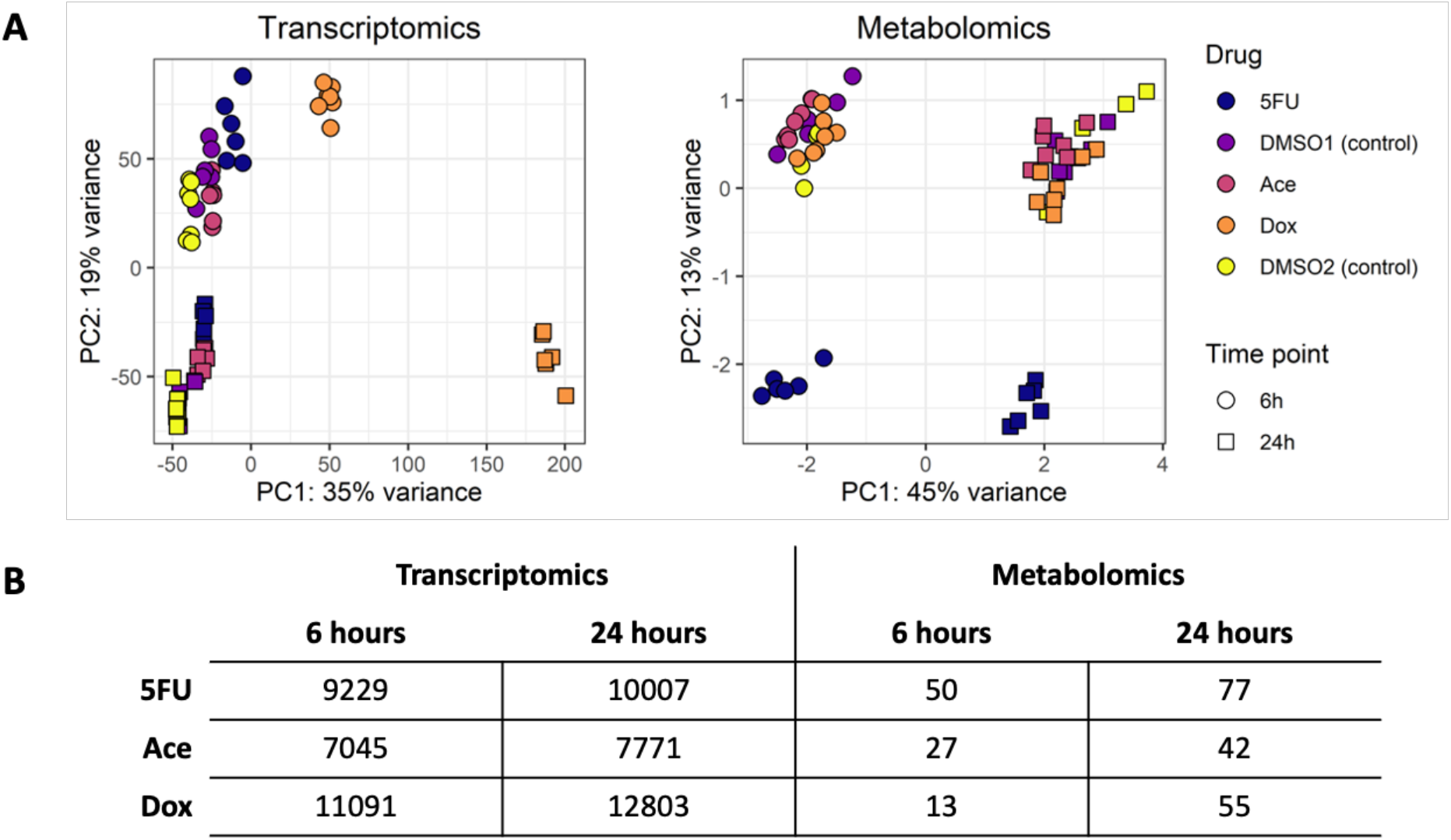
PCA of transcript counts (A) and scaled metabolite abundances (B) for the three compounds at 6 and 24 hours. (A) PCA of transcript counts separate by time and treatment condition, specifically the Dox treatment. (B) PCA of scaled metabolite abundances separate by time and treatment condition, specifically the 5FU treatment. (C) Quantification of the number of differentially expressed genes and differentially changed metabolites for each condition. DEGs were genes with an FDR < 0.01 and differential metabolites were metabolites with an FDR < 0.1.

A PCA of log-scaled and median-centered metabolite abundances separates in the first principal component by time and in the second principal component by the 5FU treatment (Figure 2B). As with the transcriptomics data, the clear separation by time point suggests a potential adaption of the primary cells to *in vitro* culture conditions. A bi-plot (Supplemental Figure 3B) of the top 10 metabolites responsible for the PCA separation identifies erythritol, a derivative of glucose metabolism (Schlicker et al, 2019) and ethylmalonate, a branched chain fatty acid, for primary separation in the first principal component, suggesting a change in glucose and fatty acid metabolism over time. The second principal component separates by the 5FU treatment. Given that 5FU’s chemotherapeutic mechanism of action is acting as an analogue for uracil and interfering with RNA synthesis (Zhang et al, 2008), it is not surprising to see clear separation in the metabolomics data for cells treated with 5FU. A bi-plot (Supplemental Figure 3B) of the top 10 metabolites responsible for the separation identifies uracil, phosphate, and 2-deoxyuridine as primarily driving separation in the second principal component, indicating changes in uracil synthesis and general metabolism as separating 5FU from the other conditions. However, in contrast to the transcriptomics results, there is no clear separation among any conditions, except for 5FU treatment. This lack of separation could result from the number and type of metabolites that were profiled or could suggest a more nuanced change between the Dox and Ace conditions and their respective controls. Finally, the number of differentially expressed genes (DEGs) and differentially changed metabolites were quantified for each treatment and time point (Figure 2C). As was expect ed, from the PCAs, the Dox condition has the largest number of DEGs at both 6 and 24-hours while the 5FU condition has the largest number of differentially changed metabolites at the 6 and 24-hour conditions.

### Gene enrichment and metabolomics data identify common signatures of toxicity but cannot readily identify mechanisms of cardiotoxicity

The large number of DEGs for each condition necessitates an enrichment approach to identify changed pathways that are shared across conditions and time points and could therefore be suggestive of a common mechanism of cardiotoxicity. Pathway enrichment analysis was performed using the 50 Hallmark gene sets defined in the MSigDB (Liberzon et al, 2016; Yu et al, 2012) (Figure 3A). Given the large number of DEGs, it was necessary to use both a lower FDR cutoff to define differential expression and a higher p-value cutoff for enriched gene sets; genes were considered differentially expressed with an FDR < 0.01 and gene sets were enriched with a BH-adjusted p-value < 0.1. Consistent with their known mechanisms of chemotherapeutic efficacy, the 5FU and Dox conditions are enriched for genes related to DNA repair at both the 6 and 24-hour timepoint. As with the PCA data, the p53 pathway is enriched for the Dox condition at both 6 and 24 hours as well as enrichment in at least one timepoint for the other two conditions. Finally, genes related to oxidative phosphorylation are enriched, particularly at 6 hours, for both 5FU and Dox, consistent with changes in cellular metabolism.

**Figure 3.**
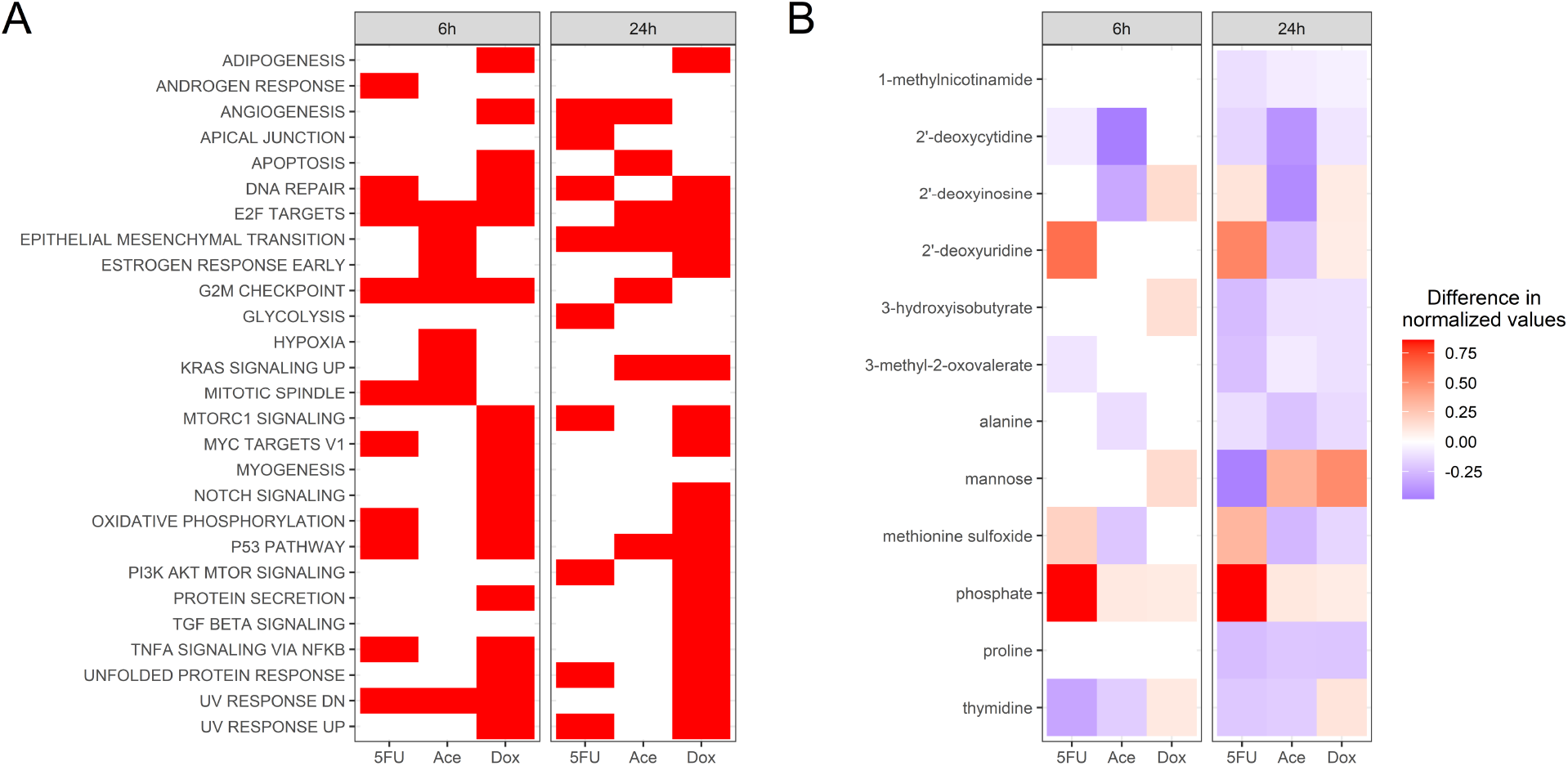
Identifying biomarkers of toxicity from the transcriptomics and metabolomics data sets. (A) Enrichment analysis for genes with an FDR < 0.01 using the Hallmark gene sets from the Molecular Signatures database. A red box indicates enrichment with a p-value < 0.1. Only gene sets that were enriched in at least condition are displayed here. (B) Metabolites that were identified to be significantly changed in production between the treatment and control (Mann-Whitney, FDR < 0.1). The color represents the mean difference in normalized metabolites between the treatment and control, where a negative value indicates decreased production and a positive value indicates increased production.

Two pathways are enriched across all conditions at 6 hours, the E2F targets and G2M checkpoint, both of which are pathways involved in the cell cycle. Only one pathway is enriched across all three conditions at 24 hours, the epithelial to mesenchymal transition (EMT) pathway.

Next, metabolites were identified that were significantly changed across all conditions within a timepoint (Figure 3B). We see increased production of phosphate for all three compounds, suggesting a significant change in metabolism for the chosen concentrations, consistent with the measured decrease in metabolic activity at the chosen concentrations (Figure 1). Further, there was a consistent increase in production of 2’-deoxyinosine for 5FU and Dox at the 24-hour timepoint, consistent with a response to reactive oxygen species (ROS) stress (Lee et al, 2015) as well as increased production of 2’-deoxyuridine in the both the 5FU and Dox condition, consistent as a by-product of DNA damage and uracil metabolism (Evans et al, 2010; Hagenkork et al, 2017). Metabolites were both (a) differentially consumed between conditions when compared to blank media controls (Supplemental Figure 4A) and (b) produced or consumed between conditions when compares to blank media controls (Supplemental Figure B). Of note is uracil; in the 5FU treatment condition at both timepoints, uracil was consumed more in the treatment compared to the control condition but in the Dox, Ace, and control conditions at 24 hours, uracil was produced. A metabolite that had a measured change in production between treatment and control (Figure 3B) can serve as potential biomarkers of *in vitro* cardiotoxicity; however, a specific mechanism for the change in production, and its relationship to changes in consumption, is difficult to assess. Further, there are no available methods for connecting the DEGs and differentially changed metabolites to identify metabolic adaptions during cardiotoxicity and potential genes driving those adaptations. Metabolic network reconstructions provide an opportunity to connect the measured changes in the transcriptome with the measured changes in the metabolome to identify these potential metabolic adaptations during cardiotoxicity.

### Reconstruction of a rat-specific heart GENRE from an existing human-specific heart GENRE

Paired GENREs of human (*iHsa*) and rat (*iRno*) metabolism have been published, which capture species-specific metabolic functions and species-specific gene-protein-reaction (GPR) rules (Blais et al, 2017). However, integrating the data collected here requires a rat, heart-specific metabolic model, which have not been published to date. A human-specific heart GENRE, *iCardio* (Dougherty et al, 2020), was recently published that was built from the paired human and rat GENREs, *iHsa* and *iRno. iCardio* was built by integrating tissue-specific protein expression data available in the Human Protein Atlas (HPA) (v18.proteinstlas.org; Ulhen et al, 2015) with the *iHsa* model followed by manual curation using pre-defined metabolic tasks. Given that *iHsa* and *iRno* were generated in parallel, the reactions included in the human-specific heart model, *iCardio*, directly map to a rat-specific heart model using the common reaction identifiers. After including these reactions, 13 rat-specific reactions were added from the general rat model, *iRno*, that had literature evidence for expression in the heart or were necessary for metabolic functions (Supplemental File 1).

Next, the metabolomics data was used to identify metabolites that were measured in the collected dataset and also included in the metabolic model. 121 metabolites mapped between the metabolomics dataset and metabolites in the model. 75 of these metabolites had associated exchange reactions in the general rat model, indicating that the metabolite could be either consumed or produced in the model. From these exchange reactions, 37 were added to the rat-specific heart model from the general model to ensure that constraints could be placed for either production or consumption. Manual curation is often a time-intensive process and here we demonstrate the value of metabolomics data in identifying new potential reactions that should be added to tissue-specific models of metabolism.

Before integrating the transcriptomics with the model, we first identified the number of DEGs and differentially changed metabolites that the new rat-specific heart model captures (Supplemental Figure 5A). The model captures ∼10% of the DEGs across all treatments and time points and between 40-65% of the differentially changed metabolites. Next, in order to confirm that the metabolic genes captured in the model are still capturing the underlying variability in the data (Figure 2A), PCA was used to identify the largest sources of variability in the metabolic genes represented in the model (Supplemental Figure 5B). As with the previous PCA (Figure 2), there is a similar spread in the data. Using a similar enrichment approach as before, both the glycolysis and hypoxia Hallmark pathways, among others, were enriched in the top 100 genes separating the first principal component, suggesting that a shift in glycolysis may have a unique role in doxorubicin toxicity. For the second principal component, the Hallmark pathways of cholesterol homeostasis, glycolysis, oxidative phosphorylation, and fatty acid metabolism, among others, were enriched. Again, this suggests the expected shift from the fetal gene program present in neonatal cells, from primarily glycolysis to fatty acid metabolism as a source of ATP. However, as with the previous results, it is unclear from the enrichment analysis alone what the direction of change is, i.e. from the fetal gene program as is seen maturing cardiomyocytes or towards the fetal gene program which has been noted in diseased cardiac status, such as heart failure (Taegtmeyer et al, 2010). Nonetheless, the model is able to capture metabolic adaptations in response to or as a result of drug treatment. However, as with the previous analysis, general pathway level changes do not yield insight into potential mechanisms of cardiotoxicity.

### *Integrating transcriptomics data with a rat-specific, heart GENRE predicts novel metabolic functions altered in* in vitro *cardiotoxicity*

Gene enrichment analyses can be helpful in identifying broad changes in a data set. However, as noted above, it can be difficult to the relationship between DEGs and metabolic shifts, even with measured metabolomics data. Here, the TIDEs approach (Dougherty et al, 2020) was used to identify metabolic functions that are associated with a significant change in gene expression using the rat-specific heart GENRE (Figure 4, Supplemental File 1). The TIDEs approach uses the complex GPR rules in a metabolic model to assign weights based on the log fold change of DEGs to individual reactions necessary to complete a metabolic task, such as production of ATP from glucose. Metabolic tasks were identified that are significantly associated with either increased or decreased gene expression in each drug condition. Statistical significance is calculated by randomizing log fold change values to create a distribution of metabolic task scores to assign significance to the original score. Using this approach, metabolic functions that are significantly associated with changes in gene expression are used to identify metabolic adaptations that are a potential cause or result of cardiotoxicity.

**Figure 4.**
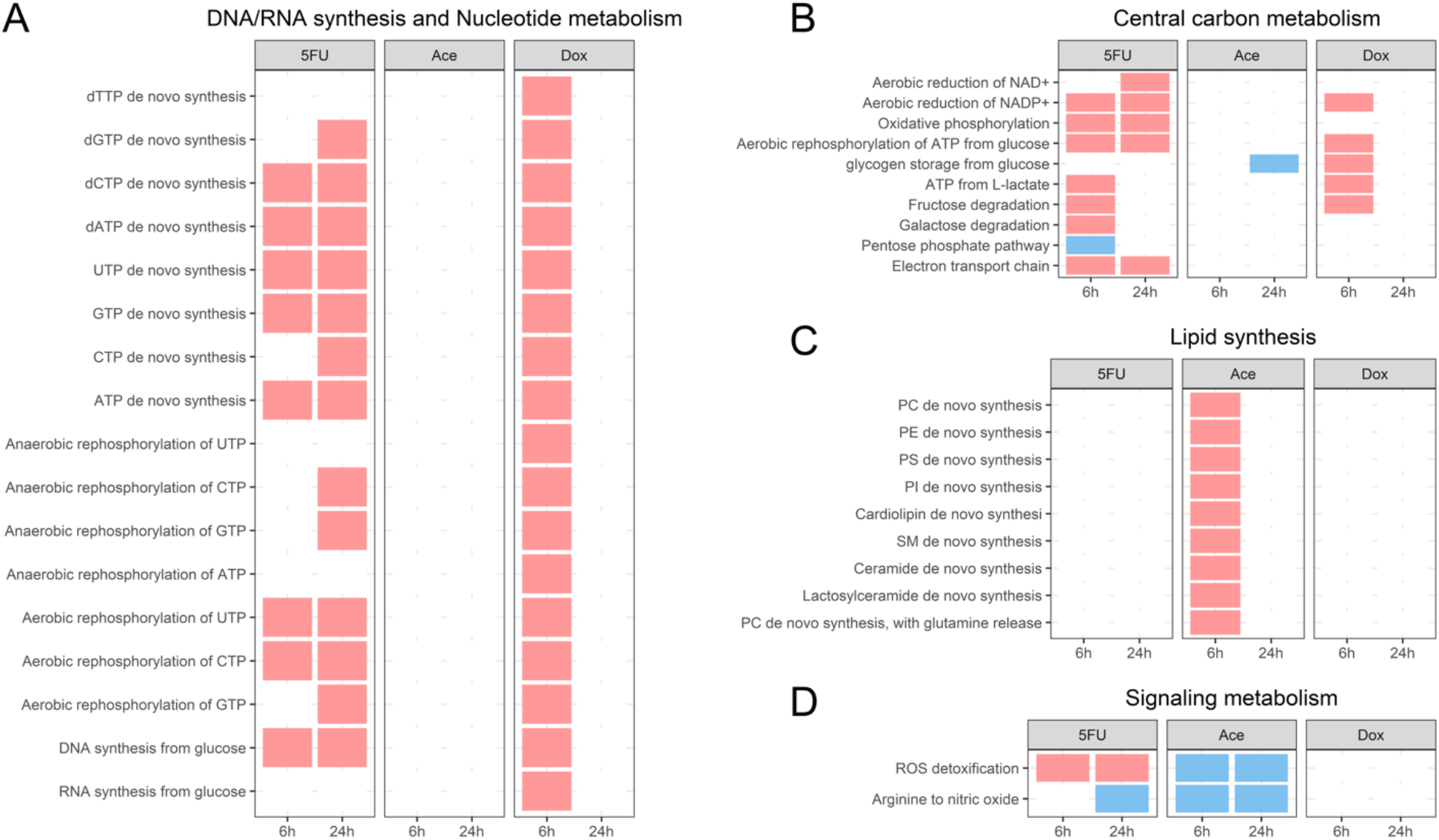
TIDEs analysis reveals distinct changes in metabolic function in response to compounds. A red box indicates significantly higher associated gene expression and a blue box indicates significantly lower associated gene expression. (A) Metabolic tasks related to DNA and RNA synthesis or nucleotide metabolism. (B) Metabolic tasks related to central carbon metabolism. (C) Metabolic tasks related to lipid membrane synthesis. (D) Metabolic tasks related to signaling metabolism.

The log fold change of metabolic genes with an FDR < 0.01 were overlayed onto the rat-specific heart GENRE (Methods, Supplemental File 1). Metabolic tasks with a p-value < 0.1 were identified as differentially changed. All metabolic tasks related to DNA/RNA synthesis and nucleotide metabolism are increased in the Dox condition at 6 hours. In contrast, 5FU has metabolic tasks related for production of nucleotides and nucleoside triphosphates (NTPs) increased at both 6 and 24 hours, except for anaerobic rephosphorylation of NTPs at 6 hours. Together, changes in both NTP and dNTP metabolism suggest that both 5FU and Dox have an effect on both RNA and DNA synthesis, which could serve as a potential metabolic stress on the cell.

Next, given that the Hallmark pathway of oxidative phosphorylation was enriched from the Hallmark gene set (Figure 3A), we examined changes in metabolic tasks related to central carbon metabolism (Figure 4B). A significant number of tasks were changed for the 5FU condition at both timepoints, indicating an overall strong metabolic shift in response to 5FU. The decrease of the metabolic task for the pentose phosphate pathway suggests that metabolic intermediates may be diverted to serve the increased metabolic demands induced by the 5FU treatment. As with the previous set of metabolic tasks, Dox is associated with significant changes in central carbon metabolism at 6 hours but no significant changes at 24 hours.

Given that Ace is not a known cardiotoxic compound, we next identified that metabolic tasks related to lipid membrane synthesis were uniquely increased in the Ace condition at 6 hours (Figure 4C). Particularly, metabolic tasks were upregulated for a variety of phospholipids, suggesting a unique metabolic adaptation in response to Ace treatment. Of particular interest, the metabolic task for cardiolipin synthesis, a key component of the mitochondrial metabolism, is significantly increased. One mechanisms of Ace hepatoxicity is associated significant lipid peroxidation in response to increased oxidative stress (Jaeschke et al, 2018), suggesting a potential shared mechanism between hepatoxicity and cardiotoxicity.

Finally, metabolic tasks that were significantly changed in signaling metabolism were identified (Figure 4D). Here, the synthesis of nitric oxide from arginine was decreased for three of the six conditions as well as differences in ROS detoxification between 5FU and Ace. Given the known role of Ace ROS production in Ace hepatotoxicity (Jaeschke et al, 2018), it is interesting to see decreased gene expression for the metabolic task of ROS detoxification and the pentose phosphate pathway. Decreased synthesis of NO has been noted in heart failure (Dougherty et al, 2020; Li et al, 2020; Massion et al, 2003), suggesting a shared metabolic marker of heart dysfunction.

Across all conditions and time points, the fewest significant metabolic tasks were seen in the Dox 24-hour condition, which had the largest number of DEGs. For the metabolic task of arginine to nitric oxide and ROS detoxification, the underlying distribution of randomized task scores used to determine statistical significance was examined (Supplemental Figure 5). From these data, the Dox condition has a higher overall task score, indicating a higher overall average gene expression across reactions, but also has a wider spread for the underlying distribution of randomized task scores as a result of the large number of DEGs.

### Combined omics datasets predict novel metabolic adaptations in toxicity through integrated network-based analysis

Pathway-level analysis provides one point of view for interpreting changes in gene expression. However, pathways do not work independently, but rather, act in a coordinated effort to maintain cell function. GENREs are able to capture this relationship by determining flux through reactions in a model while meeting an objective function that represents a hypothesis for cell function. Here, the transcriptomics data was integrated with the rat-specific heart model using the RIPTiDe algorithm (Jenior et al, 2020) to determine the reactions and fluxes that met the constraints provided by our objective function. A single objective function for non-proliferative cells is hard to define; here, the objective function of ATP hydrolysis, representing the ATP generated for cardiomyocyte contraction, as well as requiring minimal synthesis of DNA and RNA, was used (Methods). Using the condition-specific transcriptomics data, condition-specific models were generated for each condition, both treatment and control groups. Details for the condition-specific models are in Supplemental Table 1.

Each condition-specific model at 24 hours had a significant correlation (p-value < 0.05) between transcript counts (TPMs) and the flux samples (Supplemental Table 1). This result indicates that the context-specific patterns of metabolism predicted with RIPTiDe are significantly correlated with the experimentally measured omics data, further supporting the validity of our predictions for the underlying biology. Non-metric multidimensional scaling (NMDS) was used to visually display the 50 flux samples for each condition in an unsupervised fashion (Supplemental Figure 7). For this approach, fluxes were plotted for the 112 reactions that are shared amongst all conditions. The NMDS plots would suggest that, as was suggested in earlier PCA plots, the DMSO controls each elicit a different metabolic response.

Supervised machine learning with random forest feature selection was used to identify the reactions that most distinguish the active metabolism of treatment from the respective control conditions. This analysis highlighted that every treatment and time point contained at least one reaction involved in central carbon metabolism as highly distinguishing, suggesting unique divergent pathways for ATP production between treatment and control conditions. Second, when aggregating treatment conditions versus the control conditions, reactions were identified that were uniquely associated with changes in the treatment conditions. Flux samples for the two reactions identified as highly distinguishing (Figure 5) demonstrate differences in pyrimidine metabolism across treatment groups. In the case of 5FU and Dox, increased flux is predicted for the formation of PEP from pyruvate (Figure 5A, negative flux), where phosphoenolpyruate (PEP) is used as an intermediate in the formation of UTP from UDP (Figure 5A, positive flux). The predicted flux for the increased production of UTP again highlights that the primary chemotherapeutic mechanisms of action for 5FU and Dox (DNA and/or RNA damage) also serves to generate increased metabolic stress. It is important to note that the same constraints were placed on the overall network structure in this case (i.e. the same level of production of DNA and RNA). However, the transcript data imposed another constraint which, in this case, identified the first reaction as different in all three treatment conditions. Looking at the transcript data (Figure 5B), the 5FU and Dox condition have elevated transcript counts for *PKM2*, one of the genes responsible for catalyzing these two reactions.

**Figure 5.**
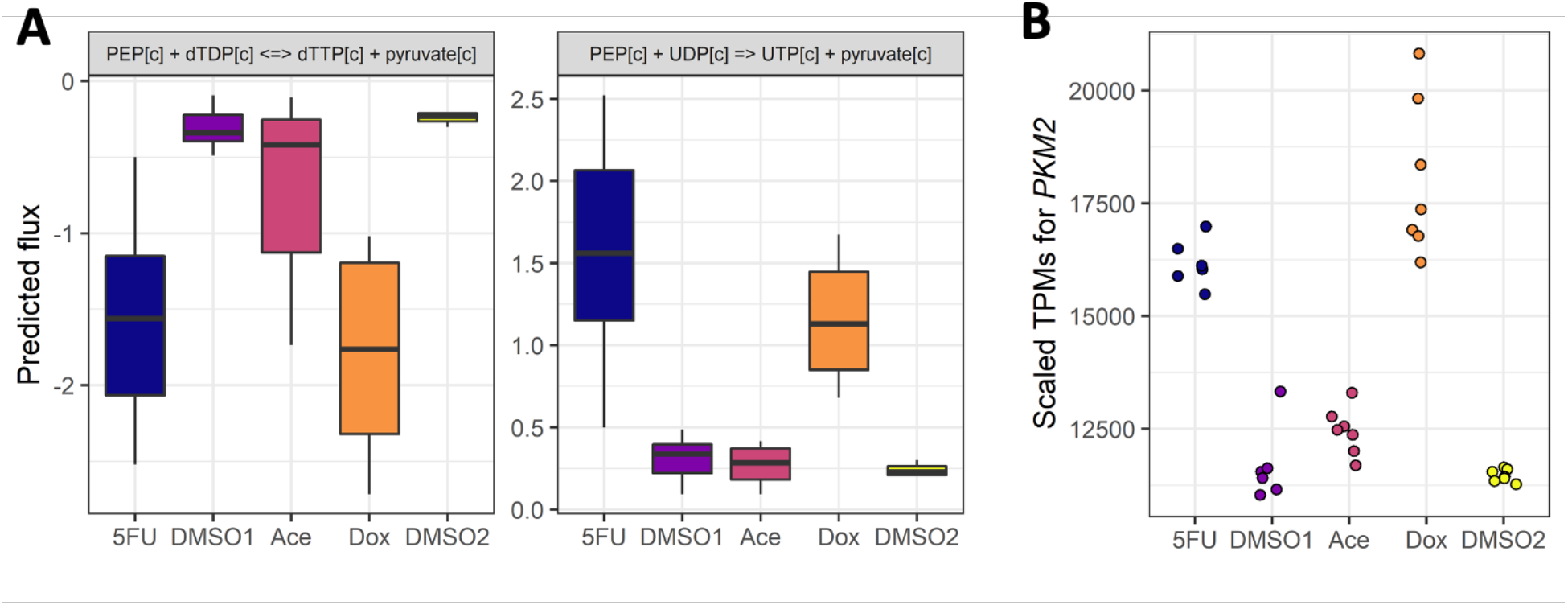
Condition-specific models integrating the metabolomics and transcriptomics 24 hour data identify unique reaction common across treatment groups that is driven by changes in transcription. (A) Distribution of predicted fluxes through two reactions that were predicted to most strongly predict the difference between the treatment and control conditions. (B) Transcript abundances for one of the genes responsible for catalyzing both the reactions in panel A.

## Discussion

Here, we present paired transcriptomics and metabolomics data characterizing *in vitro* cardiotoxicity for three compounds: 5-fluorouracil, acetaminophen, and doxorubicin. Cardiotoxic concentrations were identified as concentrations that induce a change in metabolism without a significant change in cell death for all three compounds. The data provides the first, to our knowledge, paired transcriptomic and metabolic characterization of *in vitro* cardiotoxicity. For the transcriptomic data, Dox elicits the strongest transcriptomic response, followed by time. For the metabolomics data, time elicits the strongest metabolic response, followed by the 5FU condition. In the metabolomics data, in contrast to the transcriptomic data, no difference in conditions outside the 5FU treatment was observed, indicating either a lack of sufficient coverage to identify metabolic differences or more nuanced changes in Dox and Ace and their respective controls.

The treatment was confirmed using enrichment analysis, demonstrating significant enrichment for DNA repair in both 5FU and Dox, where both chemotherapeutic mechanisms of action target DNA or RNA synthesis. Additionally, metabolic changes were seen through enrichment in oxidative phosphorylation and increased release of phosphate measured in the metabolomics data across all conditions. The only Hallmark pathway that was enriched across all conditions at 24 hours was the EMT pathway. EMT is an important regulator in the development of the heart and, more recently, has been shown to have a role in the development of cardiac fibrosis (von Gise et al, 2012). Here, enrichment in genes in the EMT pathway could suggest a role for EMT in cardiotoxicity. While pathway level and metabolite changes are helpful, there are few methods currently available that use known mechanisms to use changes at the gene level to propose potential metabolic adaptations in response to treatment.

While standard differential gene expression analyses and differential metabolite analyses are useful, models of metabolism, such as GENREs, provide an opportunity to give mechanistic insight into the relationship between changes in expression and metabolism. These approaches provide a two-fold perspective by first (a) reducing the set of DEGs and reactions down to key metabolic reactions, and (b) providing a direct relationship between a prediction (i.e. a metabolic task or flux) and gene expression. Here, a key driver of the separation seen in the gene expression data (Figure 1) is metabolically related genes (Supplemental Figure 5B), which was not identified in standard enrichment analyses. Second, we identify a metabolic reaction that serves as distinguishing between treatment and control conditions (Figure 5A) and then gene that is driving that prediction (Figure 5B).

Consistent with the enrichment analysis, integration of DEGs with the TIDEs approach specifically identified changes in metabolic tasks related to DNA/RNA synthesis but also specifically identifying the role of all nucleotides in both Dox and 5FU. In the case of Ace, the TIDEs pipeline identified increased synthesis of multiple phospholipids metabolic adaptations during cardiotoxicity. Lipid peroxidation of membrane phospholipids is one proposed mechanism of acetaminophen-induced hepatoxicity (Jaeschke et al, 2018), but has not been proposed in cardiomyocytes. Nitric oxide synthesis and ROS detoxification were identified as common metabolic functions altered across conditions. Nitric oxide synthesis is a proposed mechanism of doxorubicin-induced cardiotoxicity (Bahadir et al, 2014) but has not yet been identified for 5FU cardiotoxicity. Here, decreased NO synthesis and changes in ROS detoxification were identified as early and shared markers of *in vitro* cardiotoxicity. Finally, it is important to note the difference in number of TIDEs identified between conditions, where the Dox 24-hour condition has the lowest number of TIDEs. Given that there are the largest number of DEGs in the Dox condition, most genes that were included in the TIDEs analysis for the Dox 24-hour condition were differentially expressed, meaning that a metabolic function had to have a large gene expression signature to be identified as significant (Supplemental Figure 5). This is one weakness of the current approach. Future approaches should be work towards being independent of the number of DEGs and should instead rely on transcript counts in the treatment and control conditions to determine shifts in metabolic functions.

Although enrichment analyses are helpful in determining broad metabolic changes, it is important to view these changes in the context of the entire metabolic network. RIPTiDe provides both the reactions necessary and predicted fluxes through those reactions that satisfied the given objective function and the transcriptomics constraints. In this case, it is important to highlight that the chosen objective function, synthesis of ATP, DNA, and RNA, utilizes only central and key metabolic reactions. More complex objective functions provide the opportunity to explore additional, peripheral metabolic pathways. However, it is difficult to define appropriate objective functions for human metabolism. Therefore, for this study, we chose to focus on the core metabolic functions of ATP synthesis, representing cardiac contraction, and baseline DNA and RNA synthesis.

Random forest variable selection identified reactions that were shared among individual conditions but whose flux separated between conditions. These reactions represent changes in flux that are necessary either for ATP production, metabolite production, or DNA/RNA synthesis that differ between condition, indicating divergent pathways of flux. Across all three treatment groups, reactions for central carbon metabolism were identified as significantly different between conditions, indicating a baseline divergent flux for ATP production. When comparing all treatment conditions to the control conditions, reactions related to the formation of PEP were significantly different, where PEP serves as an intermediate in the synthesis of UTP for the 5FU and Dox conditions. These predicted fluxes are confirmed when looking at the transcript counts (Figure 5B) for one of the enzymes (*PKM2*) that catalyzes this reaction. It is important to note the decreased flux seen in the formation of PEP in the Ace condition with no consumption of PEP in the reaction that is utilized for the 5FU and Dox conditions. This indicates that while a similar pathway is used for PEP synthesis, PEP is redirected to another metabolic pathway in the Ace condition, highlighting the interconnected and complex dynamics of metabolism.

Together, the paired transcriptomics and metabolomics data taken with predictions made using the rat-specific model of metabolism provide mechanistic insight that was not clear from either set of data on its own. For Dox, we identified shifts in metabolic tasks related to nucleotide metabolism, ROS detoxication, and NO synthesis, consistent with previously published hypotheses for mechanisms of toxicity (Farias et al, 2017; Deidda et al, 2016), as well as shifts in nucleotide metabolism as a potential additional metabolic stress contributing to cardiotoxicity. For 5FU, we identified shifts in metabolic tasks related to nucleotide metabolism, ROS detoxification, and NO synthesis, and, as with Dox, shifts in nucleotide metabolism suggesting increased metabolic stress from the chemotherapeutic mechanism of action of 5FU. Finally, for Ace, we identified shifts in metabolic tasks related to lipid synthesis and a role for PEP synthesis as a metabolic adaptation in response to toxicity.

Future work is necessary to trace pathway fluxes to determine how fluxes through these individual reactions influence other parts of metabolism. The collected metabolomics data also serve as an additional set of constraints to place when integrating the transcriptomics data with GENREs. However, more work needs to be done in developing methods to identify the relationship between a metabolic constraint and a change in predicted flux. In addition, the provided paired transcriptomics and metabolomics data provide a starting point for improvements to the present metabolic network reconstruction. A number of metabolites were measured as produced but could not be produced with the individual condition-specific models, either because of missing internal reactions or missing constraints. Finally, future work can explore the use of additional objective functions that replicate proposed mechanisms of toxicity, such as increased ROS production or synthesis of key cellular proteins, which may provide further explanation for the measured changes in the metabolomics data.

## Methods

### In vitro *culture conditions*

Primary neonatal rat cardiomyocytes were isolated and cultured according to previously published protocols (Ryall et al, 2014). After the initial plating, cells were maintained for ∼36 hours in plating media containing low glucose DMEM and M199 supplemented with L-glutamine, Penicillin-Streptomycin, 10% horse serum and 5% FBS. Cells were serum starved overnight (∼12 hours) before running experiments in serum free, ITSS-supplemented plating media. Cells were observed to beat spontaneously within 24-48 hours after isolation, confirming metabolic activity and functionality.

For experiments to determine optimal drug concentrations, cells were seeded in 96 well plates at a density of 100k cell/well. The initial range of concentrations used for 5FU, Dox, and Ace (Tocris) were selected based off previous studies for Dox (Lamberti et al, 2012), 5-FU (Lamberti et al, 2012; Lamberti et al, 2014), and Ace (Jin et al, 2012). Drug stocks were prepared according to manufacturer’s instructions using sterile DMSO and were diluted in plating media before treatment. Concentrations that induced cardiotoxicity were determined using parallel measures of cell death and cell reducing potential (10 mM 5FU, 2.5 mM Ace, 1.25 µM Dox). Cell death was determined as the number of propidium iodide (PI) positive cells divided by the total number of Hoescht positive cells. Fluorescence data from treated cells was background corrected using blank wells before using CellProfiler (McQuin et al, 2018) to segment nuclei and measure fluorescence intensity for both PI and Hoescht. In the case of doxorubicin, which is fluorescent at overlapping wavelengths with PI, wells containing the respective concentrations of doxorubicin were used for background subtraction. Measures were aggregated from four fields of view for each drug concentration. Cell viability, which measures cell reducing potential and thus cell metabolism, was measured using the RealTime-Glo MT Cell Viability kit (Promega, Catalog #G9711). Both measures were repeated for three separate wells, representing three technical replicates for each condition, as well as on three separate days using different primary cell isolations, representing three biological replicates for each condition. Statistical significance was calculated using Dunnet’s t-test (Hothorn et al, 2008) which accounts for the dose-dependent nature of the data. A p-value < 0.05 was considered statistically significant.

Oxygen Consumption Rate (OCR) for Mitochondrial Stress Test (MST) assay was measured using a Seahorse XF24 Extracellular Flux Analyzer using previously published methods (Nagdas et al, 2019). Primary rat neonatal cardiomyocytes were plated on Seahorse assay plates and cultured according to the protocol described above. MST media was an unbuffered, phenol-red free, serum-free media from above that was adjusted to a pH of 7.4 and filter-sterilized before use. For the MST assay, oligomycin, BAM15 (Cayman), and Rotenone and Antimycin A (Sigma) were injected to final concentrations of 1 µM, 10 µM, 1 µM and 2 µM respectively. The OCR for each measure was taken as the first measurement after injection with the inhibitor or the first measurement in the case of baseline. ATP production was defined as the OCR at baseline minus the OCR after the oligomycin injection. OCR for each well was normalized to well cell numbers which were collected using a PI/Hoescht stain prior to the assay.

### RNA isolation, sequencing, and analysis

For the paired transcriptomics and metabolomics data, hearts were harvested in parallel from three litters of rats on the same day. After parallel digestion, cells were mixed from all isolations before plating in 12-well plates at a density of 1.2 million cells/well. For reference, two separate DMSO controls were run (DMSO1 at 1% DMSO for the 5FU condition; DMSO2 at 0.25% DMSO for the Ace and Dox conditions). Primary rat neonatal cardiomyocytes were exposed to the chemicals mentioned above at the chosen concentrations for either 6 or 24 hours. After exposure, as has been done in previous studies (Rawls et al, 2018; Rawls et al, 2020), the cells were lysed with Trizol to begin RNA extraction. Cell lysates were mixed with chloroform and spun in phase-lock gel tubes inside a cold room and the upper phase was then decanted into new tubes. Isopropanol and glycogen were added to the mixture, incubated overnight at -20C and spun again resulting in an RNA pellet, which was washed with 75% ethanol twice. DNA was removed using the TURBO DNA-free kit (Invitrogen, #AM1907) and then RNA was quantified using the QuBit RNA Broad Range detection kit (Invitrogen, #Q10210). RNA was sent to GeneWiz (https://www.genewiz.com/en) for PolyA selection, library construction, and sequencing. RNA was sequenced using 2×150bp paired-end (PE) readings and fastq files were generated. Kallisto v 0.46.0 (Bray et al. 2016) was used to pseudo-align raw fastq files under default settings to the *Rattus norvegicus* Ensembl v96 transcriptome. Transcript abundances were then aggregated to the Entrez gene level in R v. 3.6.3 with the package tximport (Soneson et al. 2015). Genes with consistently low counts (< 10) across all samples were removed. Differentially expressed genes (DEGs) were determined using DESeq2 (Love et al. 2014) with a significance threshold of FDR < 0.1.

Principal component analysis (PCA) was performed using the variance stabilized gene counts (Love et al, 2014) with the prcomp function in R. Statistical significance for the separation between treatment and control groups was calculated using adonis2 function in the Vegan package in R (Dixon et al, 2003).

Following identification of DEGs, two approaches were used to identify pathways significantly changed in the data: enrichment using Hallmark gene sets from the Molecular Signatures Database (Liberzon et al, 2016) and <u>T</u>asks Inferred from <u>D</u>ifferential <u>E</u>xpression (TIDEs) for identifying differentially changed metabolic functions (Dougherty et al, 2020). For the enrichment analysis, due to the large number of DEGs, enrichment was determined using genes with an FDR < 0.01 and pathways were defined as statistically significant with a p-value < 0.1 following Benjamini-Hochberg (BH) correction. For the TIDEs analysis, the subset of genes that mapped to the rat model and that had an FDR < 0.01 were used; pathways with a p-value < 0.1 when compared to randomly shuffled DEGs were defined as statistically significant.

### Collecting and analyzing metabolomics data

As described above, primary rat neonatal cardiomyocytes were exposed to the compounds at the chosen concentrations. Before lysing the cells with Trizol, the cell supernatant was removed and sent to Metabolon for analysis (https://www.metabolon.com/). Raw area counts were obtained for 181 named metabolites. Metabolites that had greater than 60% missing values (13 named metabolites) were removed from the data set. Next, missing values were imputed as half of the minimum raw area count within a metabolite. Values were then log-scaled and mean-centered within a metabolite. The Mann-Whitney U-test was used to determine if a metabolite was produced or consumed relative to the blank media samples (n = 3) as well as differences between treatment and control conditions. Metabolites were considered to be significantly changed if the p-value < 0.1 following Benjamini-Hochberg correction.

### *Building a rat cardiomyocyte-specific metabolic model from the human cardiomyocyte-specific metabolic model*, iCardio

The previously published human heart metabolic model, *iCardio* (Dougherty et al, 2020), was used to build a rat-specific heart metabolic model to contextualize changes in metabolites and DEGs. All reactions that were included in the human model were included in the rat model. The 6 updates that were made in the human model (Dougherty et al, 2020) were also made to the general rat model. Since each model contains species-specific reactions, each of the 23 rat-specific reactions were manually curated to determine if they should be included in the heart; 13 were included based on literature evidence (Supplemental File 1).

Further curation was necessary based on metabolites that were measured to be either produced or consumed in the metabolomics data. Metabolites were mapped between the metabolomics data and the metabolic model using compound identifiers from the KEGG database. Exchange reactions, which are reactions in the model that transport a metabolite into or from the extracellular compartment, were added from the general rat model to the heart model if a metabolite was measured to be either consumed or produced relative to blank media in the metabolomics data. These reactions were necessary in order for a metabolite to be modeled as either produced or consumed. Reactions added back to the heart model from the general rat metabolic network are summarized in Supplemental File 1.

To identify shifted metabolic functions with the TIDEs approach, the developed rat-specific heart model was used with the previously published list of metabolic tasks (Dougherty et al, 2020). Only genes that mapped to the metabolic model were included in the analysis. For each metabolic gene, a weight was assigned based on the FDR where a gene with an FDR < 0.01 was assigned its log fold change as a weight and 0 otherwise. As with the previous publication, reaction weights were determined based on the weights for individual genes in the complex gene-protein-reaction (GPR) rules. Task scores for individual tasks were calculated as the average weight across reactions in that metabolic task. To establish statistical significance, the weights for each metabolic gene were randomly shuffled 1000 times and significance was determined by comparing the original metabolic task score to random data.

### Predicting reaction metabolic flux using the RIPTiDe algorithm

The RIPTiDe algorithm (Jenior et al, 2020) was used to integrate the transcriptomics data to identify the most likely flux distributions in the rat-specific heart model network for each of the 10 conditions, including both treatment and control groups at both time points. RIPTiDe identifies possible optimal flux distributions in a metabolic network given the cellular investments indicated by the transcriptomic abundance data. The media composition and metabolomics data were used to place constraints on metabolite consumption for each condition-specific model. Here, the cellular objective function was defined to be ATP hydrolysis, with an upper bound of 100 units of flux, and production of 1 unit of RNA and DNA, representing general cell maintenance. The exchange reactions for consumed metabolites were given a lower bound of - 10, representing a theoretical overabundance of each metabolite for the given objective. These constraints allow for metabolites to be consumed at a relatively high flux. Finally, the upper bound of internal reactions was set as 10e6 to ensure that internal fluxes were not constraining the solution space.

After applying these condition-specific constraints, condition-specific models were generated by integrating the median transcripts per million (TPM) for each gene within a condition using the RIPTiDe algorithm (Jenior et al, 2020). The RIPTiDe algorithm was run with a minimum fraction of 90% of the objective to ensure that differences in ATP flux were not the main determinants of differences between the condition-specific models. Each condition-specific model was flux-sampled 50 times to obtain a range of possible flux distributions that satisfied the pFBA assumption.

Following RIPTiDe analysis, flux samples for each condition were analyzed to identify both unique reactions for each condition and reactions whose flux separated between conditions (Jenior et al, 2020). Non-metric multidimensional scaling (NMDS) ordination of Bray-Curtis distances between flux samples, calculated using the Vegan package in R (Dixon et al, 2003), was used to visualize differences for reactions that were shared between all conditions. Finally, random forest feature selection (Calle et al, 2011) was used to determine the reactions whose fluxes most separated between each treatment and control group.

Code to reproduce this analysis is available at (https://github.com/BonnieDougherty/Cardiotoxicity). Faw FASTQ files are available on GEO166957.

## Acknowledgements

Support for this project was provided by the United States Department of Defense (W81XWH-14-C-0054 to JP), the National Institutes of Health (NIH grant HL137755 to JS), and the National Science Foundation Graduate Research Fellowship Program (awarded to BD). The opinions and assertions contained herein are the private views of the authors and are not to be construed as the official or as reflecting the views of the U.S. Army, the U.S. Department of Defense, or the Henry M. Jackson Foundation for the Advancement of Military Medicine, Inc. (HJF). This manuscript has been approved for public release with unlimited distribution. We would like to thank Bethany Wissman for assisting with the isolation of the primary rat neonatal cardiomyocytes and Laura Dunphy for her feedback on the manuscript.

## Author Contributions

BD, GK, AW, and JP conceived the study. BD, GK, BC, and SN performed experiments. BD performed the computational modeling and data analysis. BD wrote the initial draft of the manuscript. BD, KR, MJ, BC, SN, JS, GK, AW, and JP edited and wrote the final manuscript.

## Figures

**Supplemental Figure 1.**
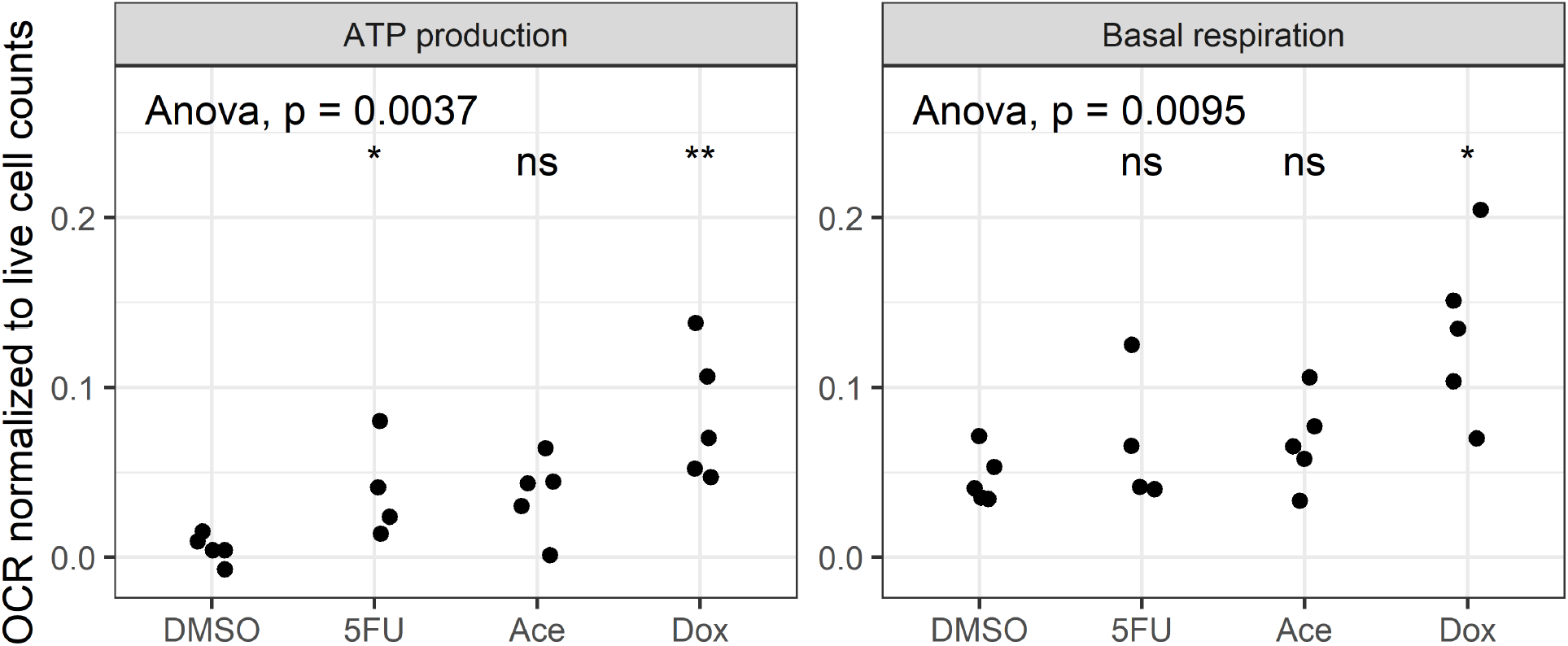
Confirming changes in measures of cellular respiration for the chosen Dox concentration using the MST assay. The MST assay was performed using the chosen concentrations for each compound. Measures of respiration where there was a statistically significant difference across groups (ANOVA, p-value < 0.05) were followed by treatment vs control comparisons using the Wilcox rank-sum test for differences where ns is not significant, ^*^ is P-value < 0.05, and ^**^ is p-value < 0.01. There were 5 experimental replicates per condition, except in the case of 5FU where one well did not respond.

**Supplemental Figure 2.**
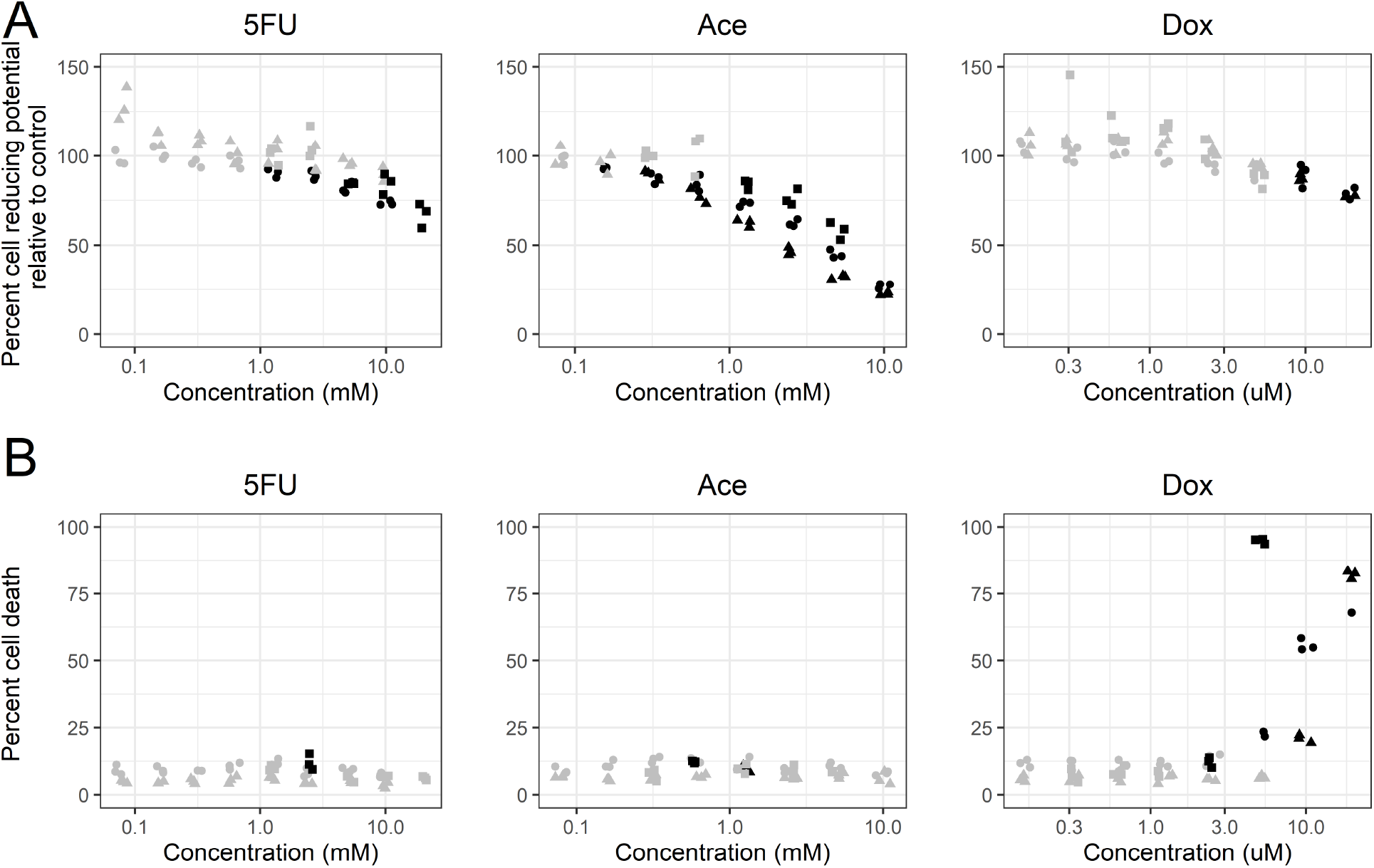
Cell reducing potential and cell death measures following 6 hours of exposure to compounds. Shapes indicate different cell isolations. Black dots indicate a statistically significant change from the control condition (p-value < 0.05) calculated using Dunntt’s test. Black boxes indicate the chosen concentrations for cardiotoxicity characterization. (A) Percent cell viability for a range of concentrations of treatment following 6 hours of exposure. (B) Percent cell death measured using a Hoescht/PI stain.

**Supplemental Figure 3.**
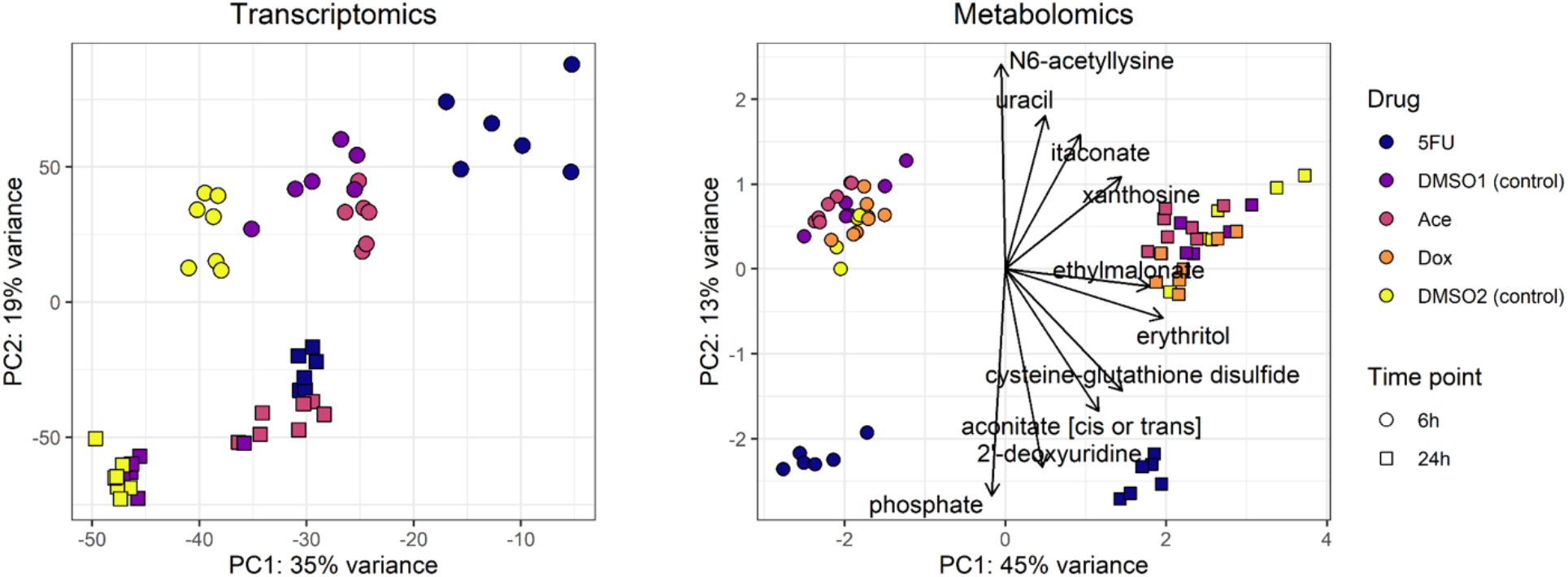
Additional PCA plots demonstrate (A) separation of treatment vs control groups and (B) the top 10 metabolites separating the PCA of the scaled metabolite abundances. (A) PCA with the Dox samples removed demonstrates separation between treated and control samples at both 6 and 24 hours. (B) The top 10 metabolites show separation in both the first and second principal component. For the first principal component, ethylmalonate and erythritol have a strong influence, suggesting a phenotypic switch between fatty acid and glucose utilization over time, although the direction is unclear. For the second principal component, phosphate and uracil separate the 5FU condition.

**Supplemental Figure 4.**
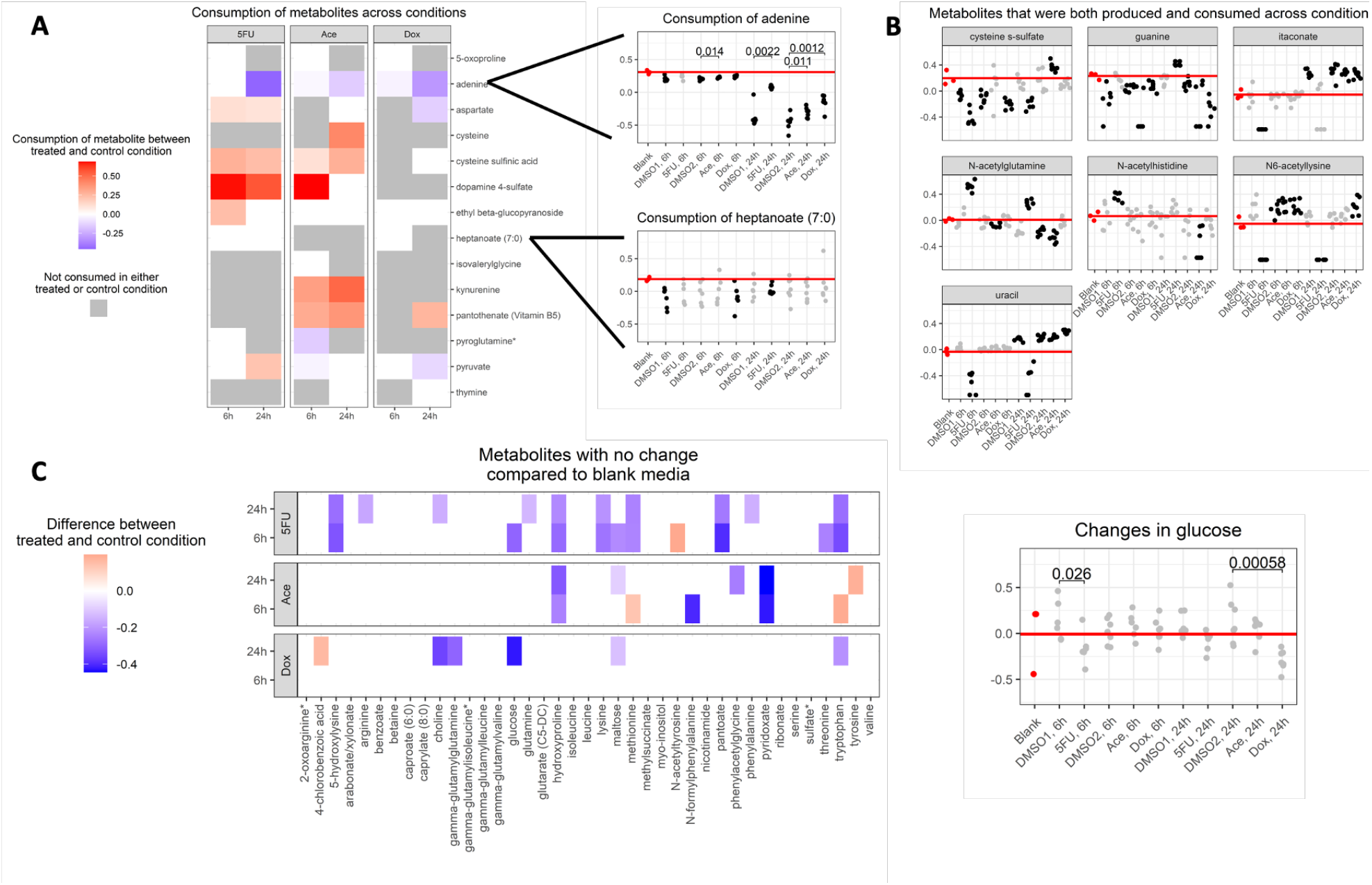
Metabolomics data shows differential consumption and production of metabolites between blank media and control/treated samples. (A) Metabolites that were measured to only be consumed across both drug treated and control conditions when compared to blank media. Here, red indicates a metabolite that was consumed more in the treatment condition than the control condition, blue indicates a metabolite that was consumed less in the treatment condition than the control condition, white indicates a metabolite that was consumed in either/both the treatment and control conditions but there was no difference in consumption and grey indicates a metabolite that was not consumed in either the treatment or control condition when compared to blank media. An example is shown for a metabolite that was more consumed between treated and control conditions (adenine) and a metabolite (hepatonate) that was consumed in a treated/control condition but there was no difference between the treated/control conditions. Black dots indicate a condition where there was a significant change in the metabolite when compared with blank media (Mann Whitney U-test, FDR < 0.1). P-values are shown for comparisons that were significant between a treatment and it’s respective control. (B) Metabolites that were measured to be both produced and consumed across conditions. Here, the red line indicates the mean value for the blank media samples. Black dots indicate conditions where there was a significant change when compared to blank media (Mann Whitney U-test, FDR < 0.1). A black dot above the red line indicates that a metabolite was produced where as a black dot below the black line indicates a metabolite that was consumed. (C) Metabolites that were not significantly changed from blank media samples but were measured to change between treatment and control samples. Here, red indicates that a metabolite was present at a higher level in the treated group compared to the control group where as blue indicates that a metabolite that present at a lower level compared to the treated group. Given that there is no significant change with respect to the blank media, directionality of change (i.e. consumption or production) cannot be determined.

**Supplemental Figure 5.**
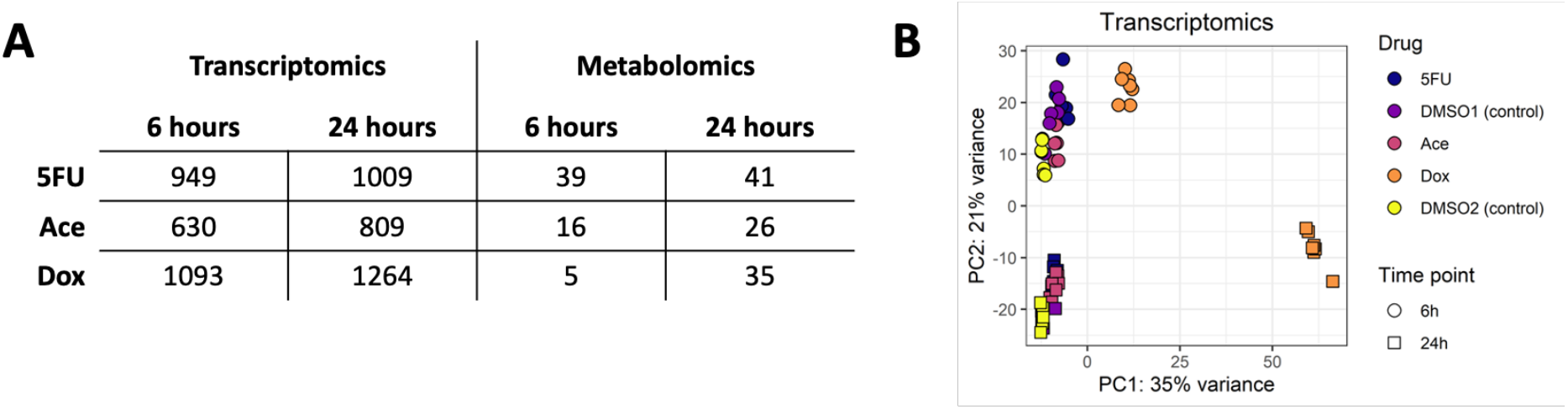
The rat-specific heart model captures changes in DEGs and metabolomics. (A) The number of DEGs (FDR < 0.1) and differentially changed metabolites (FDR < 0.1) that map to the rat-specific metabolic mode. (B) A PCA of the normalized gene counts that map back to the rat-specific heart model demonstrate clear separation, similar to the PCA of all gene counts (Figure 2), confirming that metabolism has a large determinant in separating conditions.

**Supplemental Figure 6.**
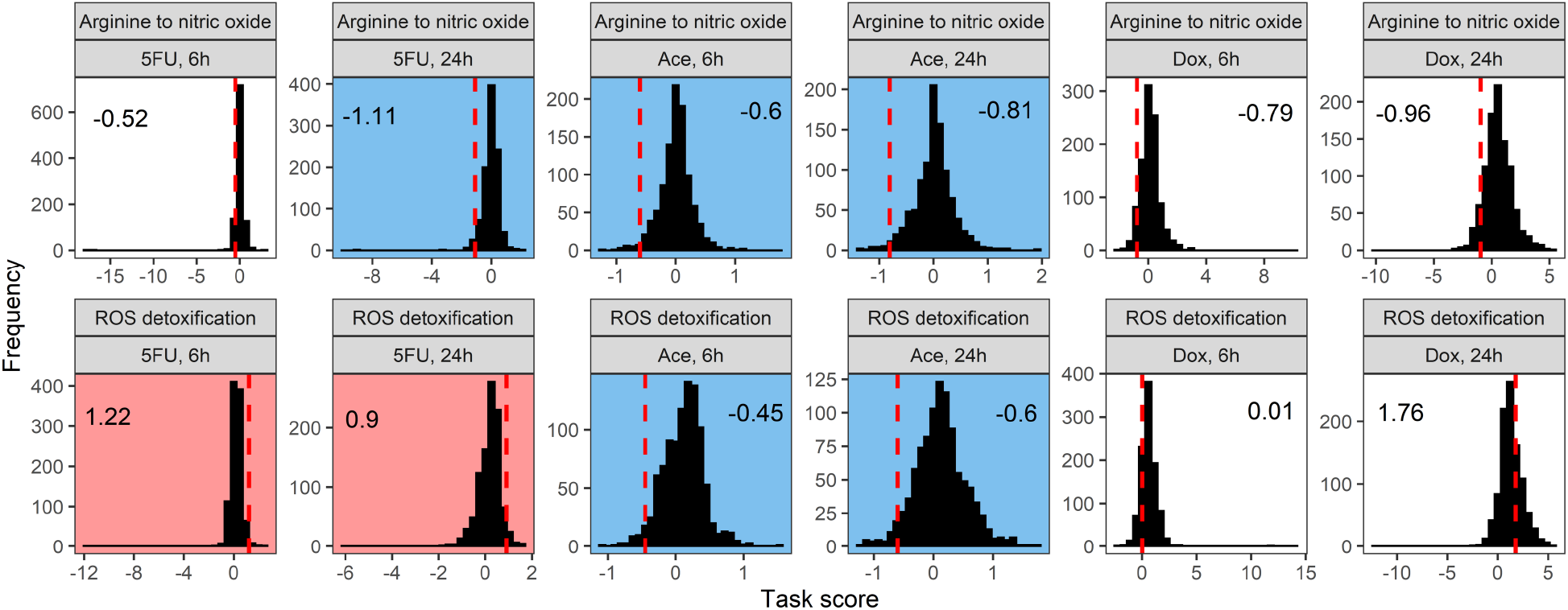
Distribution of random task scores for the metabolic tasks for arginine to nitric oxide and ROS detoxification demonstrate the underlying distribution of DEGs. A red background indicates a metabolic task associated with a significant increase in gene expression and a blue background indicates a metabolic task associated with a significant decrease in gene expression (p-value < 0.1). The red dashed line indicates the task score for the actual gene expression data whereas the black bars indicate the calculated task scores when the gene expression data is randomized. In this case, the distribution for the Dox data is significantly wider (i.e. larger range on the x-axis), indicating a larger overall absolute change in gene expression requiring a higher overall average gene expression for ROS production to be deemed significant and a lower overall average gene expression for arginine to nitric oxide to be deemed significant.

**Supplemental Figure 7.**
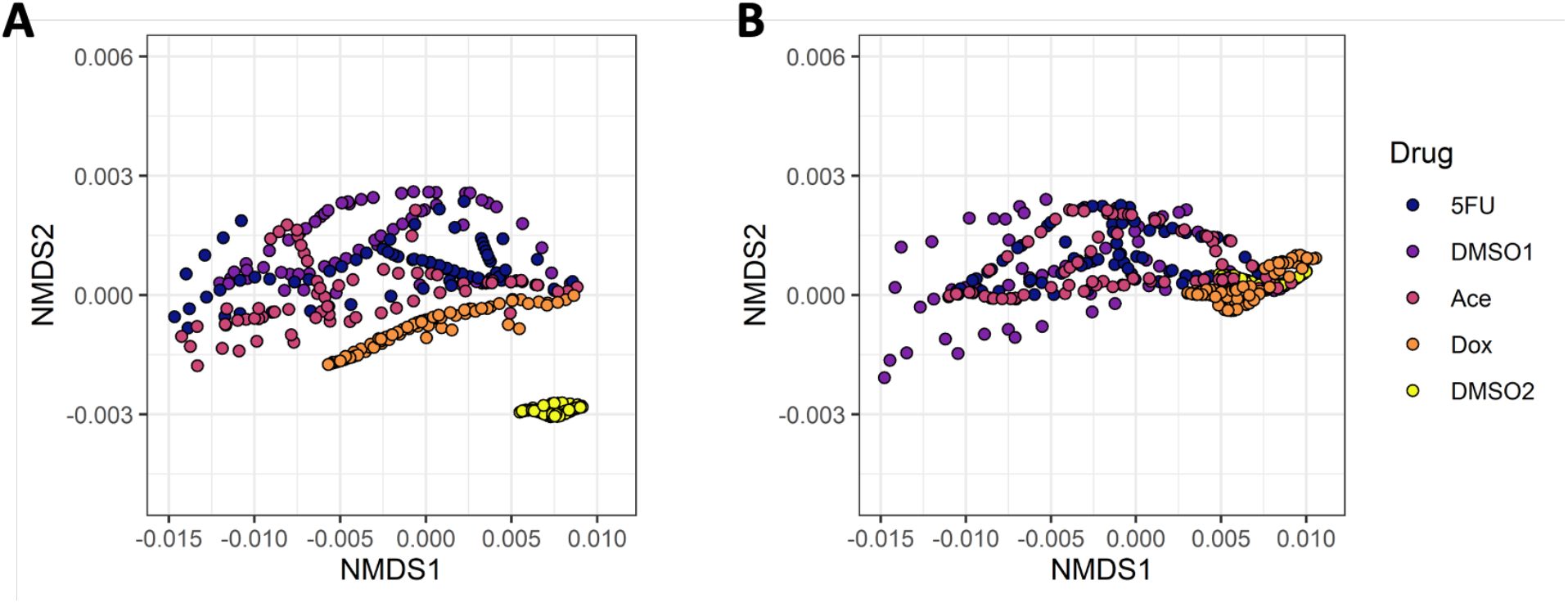
RIPTiDe models after integration with metabolomics and transcriptomics data. NMDS of the 50 flux samples for each condition at (a) 6 hours and (b) 24 hours. For each flux sample, fluxes were only taken for the 112 reactions that are shared between all conditions.

**Supplemental Table 1.**
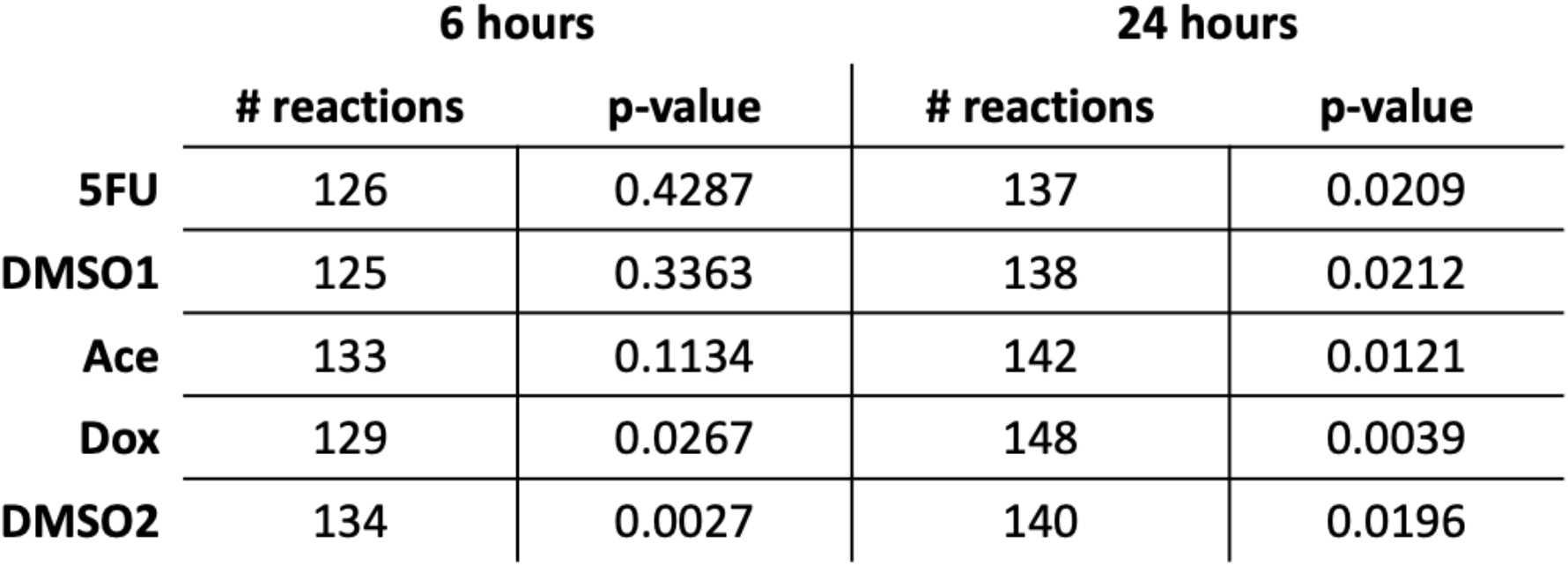
RIPTiDe models after integration with metabolomics and transcriptomics data. The number of reactions included in each model and the p-value for the Spearman correlation between reaction flux and transcript abundance.

## References

Ahuja P, Zhao P, Angelis E, Ruan H, Korge P, Olson A, Wang Y, Jin ES, Jeffrey FM, Portman M, Maclellan WR. Myc controls transcriptional regulation of cardiac metabolism and mitochondrial biogenesis in response to pathological stress in mice. J Clin Invest. 2010 May;120(5):1494–505. doi: 10.1172/JCI38331. Epub 2010 Apr 1. PMID: 20364083; PMCID: PMC2860901.

Albini A, Pennesi G, Donatelli F, Cammarota R, De Flora S, Noonan DM. Cardiotoxicity of anticancer drugs: the need for cardio-oncology and cardio-oncological prevention. J Natl Cancer Inst. 2010 Jan 6;102(1):14–25. doi: 10.1093/jnci/djp440. Epub 2009 Dec 10. PMID: 20007921; PMCID: PMC2802286.

Bahadir A, Kurucu N, Kadioğlu M, Yenilme E. The role of nitric oxide in Doxorubicin-induced cardiotoxicity: experimental study. Turk J Haematol. 2014 Mar;31(1):68–74. doi: 10.4274/Tjh.2013.0013. Epub 2014 Mar 5. PMID: 24764732; PMCID: PMC3996644.

Bauckneht M, Morbelli S, Fiz F, Ferrarazzo G, Piva R, Nieri A, Sarocchi M, Spallarossa P, Canepari ME, Arboscello E, Bellodi A, Massaia M, Gallamini A, Bruzzi P, Marini C, Sambuceti G. A Score-Based Approach to 18F-FDG PET Images as a Tool to Describe Metabolic Predictors of Myocardial Doxorubicin Susceptibility. Diagnostics (Basel). 2017 Oct 26;7(4):57. doi: 10.3390/diagnostics7040057. PMID: 29072629; PMCID: PMC5745393.

Blais EM, Rawls KD, Dougherty BV, Li ZI, Kolling GL, Ye P, Wallqvist A, Papin JA. Reconciled rat and human metabolic networks for comparative toxicogenomics and biomarker predictions. Nat Commun. 2017 Feb 8;8:14250. doi: 10.1038/ncomms14250. PMID: 28176778; PMCID: PMC5309818.

Borde C, Kand P, Basu S. Enhanced myocardial fluorodeoxyglucose uptake following Adriamycin-based therapy: Evidence of early chemotherapeutic cardiotoxicity? World J Radiol. 2012 May 28;4(5):220–3. doi: 10.4329/wjr.v4.i5.220. PMID: 22761982; PMCID: PMC3386534.

Bray NL, Pimentel H, Melsted P, Pachter L. Near-optimal probabilistic RNA-seq quantification. Nat Biotechnol. 2016 May;34(5):525-7. doi: 10.1038/nbt.3519. Epub 2016 Apr 4. Erratum in: Nat Biotechnol. 2016 Aug 9;34(8):888. PMID: 27043002.

Calle ML, Urrea V, Boulesteix AL, Malats N. AUC-RF: a new strategy for genomic profiling with random forest. Hum Hered. 2011;72(2):121–32. doi: 10.1159/000330778. Epub 2011 Oct 11. PMID: 21996641.

Deidda M, Mercurio V, Cuomo A, Noto A, Mercuro G, Cadeddu Dessalvi C. Metabolomic Perspectives in Antiblastic Cardiotoxicity and Cardioprotection. Int J Mol Sci. 2019 Oct 4;20(19):4928. doi: 10.3390/ijms20194928. PMID: 31590338; PMCID: PMC6801977.

Dixon P. VEGAN, a package of R functions for community ecology. Journal of Vegetation Science. 2003. p. 927.

Dougherty BV, Rawls KD, Kolling GL, Vinnakota KC, Wallqvist A, Papin JA. Identifying functional metabolic shifts in heart failure with the integration of omics data and a heart-specific, genome-scale model. Cell Rep. 2021 Mar 9;34(10):108836. doi: 10.1016/j.celrep.2021.108836. PMID: 33691118.

Evans MD, Saparbaev M, Cooke MS. DNA repair and the origins of urinary oxidized 2’-deoxyribonucleosides. Mutagenesis. 2010 Sep;25(5):433–42. doi: 10.1093/mutage/geq031. Epub 2010 Jun 3. PMID: 20522520.

FarÍas JG, Molina VM, Carrasco RA, Zepeda AB, Figueroa E, Letelier P, Castillo RL. Antioxidant Therapeutic Strategies for Cardiovascular Conditions Associated with Oxidative Stress. Nutrients. 2017 Sep 1;9(9):966. doi: 10.3390/nu9090966. PMID: 28862654; PMCID: PMC5622726.

Harris SL, Levine AJ. The p53 pathway: positive and negative feedback loops. Oncogene. 2005 Apr 18;24(17):2899–908. doi: 10.1038/sj.onc.1208615. PMID: 15838523.

Hothorn T, Bretz F, Westfall P. Simultaneous inference in general parametric models. Biom J. 2008 Jun;50(3):346–63. doi: 10.1002/bimj.200810425. PMID: 18481363.

Jackson T, Allard MF, Sreenan CM, Doss LK, Bishop SP, Swain JL. The c-myc proto-oncogene regulates cardiac development in transgenic mice. Mol Cell Biol. 1990 Jul;10(7):3709–16. doi: 10.1128/mcb.10.7.3709-3716.1990. PMID: 1694017; PMCID: PMC360819.

Jaeschke H, Ramachandran A. Oxidant Stress and Lipid Peroxidation in Acetaminophen Hepatotoxicity. React Oxyg Species (Apex). 2018 May;5(15):145–158. Epub 2018 May 1. PMID: 29682614; PMCID: PMC5903282.

Jenior ML, Moutinho TJ Jr, Dougherty BV, Papin JA. Transcriptome-guided parsimonious flux analysis improves predictions with metabolic networks in complex environments. PLoS Comput Biol. 2020 Apr 16;16(4):e1007099. doi: 10.1371/journal.pcbi.1007099. PMID: 32298268; PMCID: PMC7188308.

Jin SM, Park K. Acetaminophen induced cytotoxicity and altered gene expression in cultured cardiomyocytes of h(9)c(2) cells. Environ Health Toxicol. 2012;27:e2012011. doi: 10.5620/eht.2012.27.e2012011. Epub 2012 Apr 26. PMID: 22639738; PMCID: PMC3355274.

Lamberti M, Porto S, Marra M, Zappavigna S, Grimaldi A, Feola D, Pesce D, Naviglio S, Spina A, Sannolo N, Caraglia M. 5-Fluorouracil induces apoptosis in rat cardiocytes through intracellular oxidative stress. J Exp Clin Cancer Res. 2012 Jul 19;31(1):60. doi: 10.1186/1756-9966-31-60. PMID: 22812382; PMCID: PMC3461434.

Lamberti M, Porto S, Zappavigna S, Addeo E, Marra M, Miraglia N, Sannolo N, Vanacore D, Stiuso P, Caraglia M. A mechanistic study on the cardiotoxicity of 5-fluorouracil in vitro and clinical and occupational perspectives. Toxicol Lett. 2014 Jun 16;227(3):151–6. doi: 10.1016/j.toxlet.2014.03.018. Epub 2014 Apr 1. PMID: 24704391.

Lee HG, Chen Q, Wolfram JA, Richardson SL, Liner A, Siedlak SL, Zhu X, Ziats NP, Fujioka H, Felsher DW, Castellani RJ, Valencik ML, McDonald JA, Hoit BD, Lesnefsky EJ, Smith MA. Cell cycle re-entry and mitochondrial defects in myc-mediated hypertrophic cardiomyopathy and heart failure. PLoS One. 2009 Sep 25;4(9):e7172. doi: 10.1371/journal.pone.0007172. PMID: 19779629; PMCID: PMC2747003.

Li J, Kemp BA, Howell NL, Massey J, Mińczuk K, Huang Q, Chordia MD, Roy RJ, Patrie JT, Davogustto GE, Kramer CM, Epstein FH, Carey RM, Taegtmeyer H, Keller SR, Kundu BK. Metabolic Changes in Spontaneously Hypertensive Rat Hearts Precede Cardiac Dysfunction and Left Ventricular Hypertrophy. J Am Heart Assoc. 2019 Feb 19;8(4):e010926. doi: 10.1161/JAHA.118.010926. PMID: 30764689; PMCID: PMC6405673.

Li M, Parker BL, Pearson E, Hunter B, Cao J, Koay YC, Guneratne O, James DE, Yang J, Lal S, O’Sullivan JF. Core functional nodes and sex-specific pathways in human ischaemic and dilated cardiomyopathy. Nat Commun. 2020 Jun 2;11(1):2843. doi: 10.1038/s41467-020-16584-z. PMID: 32487995; PMCID: PMC7266817.

Liberzon A, Birger C, Thorvaldsdóttir H, Ghandi M, Mesirov JP, Tamayo P. The Molecular Signatures Database (MSigDB) hallmark gene set collection. Cell Syst. 2015 Dec 23;1(6):417–425. doi: 10.1016/j.cels.2015.12.004. PMID: 26771021; PMCID: PMC4707969.

Love MI, Huber W, Anders S. Moderated estimation of fold change and dispersion for RNA-seq data with DESeq2. Genome Biol. 2014;15(12):550. doi: 10.1186/s13059-014-0550-8. PMID: 25516281; PMCID: PMC4302049.

Massion PB, Feron O, Dessy C, Balligand JL. Nitric oxide and cardiac function: ten years after, and continuing. Circ Res. 2003 Sep 5;93(5):388–98. doi: 10.1161/01.RES.0000088351.58510.21. PMID: 12958142.

McQuin C, Goodman A, Chernyshev V, Kamentsky L, Cimini BA, Karhohs KW, Doan M, Ding L, Rafelski SM, Thirstrup D, Wiegraebe W, Singh S, Becker T, Caicedo JC, Carpenter AE. CellProfiler 3.0: Next-generation image processing for biology. PLoS Biol. 2018 Jul 3;16(7):e2005970. doi: 10.1371/journal.pbio.2005970. PMID: 29969450; PMCID: PMC6029841.

Nagdas S, Kashatus JA, Nascimento A, Hussain SS, Trainor RE, Pollock SR, Adair SJ, Michaels AD, Sesaki H, Stelow EB, Bauer TW, Kashatus DF. Drp1 Promotes KRas-Driven Metabolic Changes to Drive Pancreatic Tumor Growth. Cell Rep. 2019 Aug 13;28(7):1845-1859.e5. doi: 10.1016/j.celrep.2019.07.031. PMID: 31412251; PMCID: PMC6711191.

Pannala VR, Vinnakota KC, Estes SK, Trenary I, O Brien TP, Printz RL, Papin JA, Reifman J, Oyama T, Shiota M, Young JD, Wallqvist A. Genome-Scale Model-Based Identification of Metabolite Indicators for Early Detection of Kidney Toxicity. Toxicol Sci. 2020 Feb 1;173(2):293–312. doi: 10.1093/toxsci/kfz228. PMID: 31722432; PMCID: PMC8000070.

Pannala VR, Estes SK, Rahim M, Trenary I, O’Brien TP, Shiota C, Printz RL, Reifman J, Oyama T, Shiota M, Young JD, Wallqvist A. Mechanism-based identification of plasma metabolites associated with liver toxicity. Toxicology. 2020 Aug;441:152493. doi: 10.1016/j.tox.2020.152493. Epub 2020 May 30. PMID: 32479839.

Pannala VR, Vinnakota KC, Rawls KD, Estes SK, O’Brien TP, Printz RL, Papin JA, Reifman J, Shiota M, Young JD, Wallqvist A. Mechanistic identification of biofluid metabolite changes as markers of acetaminophen-induced liver toxicity in rats. Toxicol Appl Pharmacol. 2019 Jun 1;372:19–32. doi: 10.1016/j.taap.2019.04.001. Epub 2019 Apr 8. PMID: 30974156; PMCID: PMC6599641.

Rawls KD, Blais EM, Dougherty BV, Vinnakota KC, Pannala VR, Wallqvist A, Kolling GL, Papin JA. Genome-Scale Characterization of Toxicity-Induced Metabolic Alterations in Primary Hepatocytes. Toxicol Sci. 2019 Dec 1;172(2):279–291. doi: 10.1093/toxsci/kfz197. PMID: 31501904; PMCID: PMC6876259.

Rawls KD, Dougherty BV, Vinnakota KC, Pannala VR, Wallqvist A, Kolling GL, Papin JA. Predicting changes in renal metabolism after compound exposure with a genome-scale metabolic model. Toxicol Appl Pharmacol. 2021 Feb 1;412:115390. doi:10.1016/j.taap.2020.115390. Epub 2020 Dec 31. PMID: 33387578; PMCID: PMC7859602.

Ryall KA, Bezzerides VJ, Rosenzweig A, Saucerman JJ. Phenotypic screen quantifying differential regulation of cardiac myocyte hypertrophy identifies CITED4 regulation of myocyte elongation. J Mol Cell Cardiol. 2014 Jul;72:74–84. doi: 10.1016/j.yjmcc.2014.02.013. Epub 2014 Mar 5. PMID: 24613264; PMCID: PMC4078663.

Sara JD, Kaur J, Khodadadi R, Rehman M, Lobo R, Chakrabarti S, Herrmann J, Lerman A, Grothey A. 5-fluorouracil and cardiotoxicity: a review. Ther Adv Med Oncol. 2018 Jun 18;10:1758835918780140. doi: 10.1177/1758835918780140. PMID: 29977352; PMCID: PMC6024329.

Schlicker L, Szebenyi DME, Ortiz SR, Heinz A, Hiller K, Field MS. Unexpected roles for ADH1 and SORD in catalyzing the final step of erythritol biosynthesis. J Biol Chem. 2019 Nov 1;294(44):16095–16108. doi: 10.1074/jbc.RA119.009049. Epub 2019 Sep 11. PMID: 31511322; PMCID: PMC6827307.

Soneson C, Love MI, Robinson MD. Differential analyses for RNA-seq: transcript-level estimates improve gene-level inferences. F1000Res. 2015 Dec 30;4:1521. doi: 10.12688/f1000research.7563.2. PMID: 26925227; PMCID: PMC4712774.

Subramanian A, Tamayo P, Mootha VK, Mukherjee S, Ebert BL, Gillette MA, Paulovich A, Pomeroy SL, Golub TR, Lander ES, Mesirov JP. Gene set enrichment analysis: a knowledge-based approach for interpreting genome-wide expression profiles. Proc Natl Acad Sci U S A. 2005 Oct 25;102(43):15545–50. doi: 10.1073/pnas.0506580102. Epub 2005 Sep 30. PMID: 16199517; PMCID: PMC1239896.

Taegtmeyer H, Sen S, Vela D. Return to the fetal gene program: a suggested metabolic link to gene expression in the heart. Ann N Y Acad Sci. 2010 Feb;1188:191–8. doi: 10.1111/j.1749-6632.2009.05100.x. PMID: 20201903; PMCID: PMC3625436.

Taymaz-Nikerel H, Karabekmez ME, Eraslan S, Kırdar B. Doxorubicin induces an extensive transcriptional and metabolic rewiring in yeast cells. Sci Rep. 2018 Sep 12;8(1):13672. doi: 10.1038/s41598-018-31939-9. PMID: 30209405; PMCID: PMC6135803.

Uhlén M, Fagerberg L, Hallström BM, Lindskog C, Oksvold P, Mardinoglu A, Sivertsson Å, Kampf C, Sjöstedt E, Asplund A, Olsson I, Edlund K, Lundberg E, Navani S, Szigyarto CA, Odeberg J, Djureinovic D, Takanen JO, Hober S, Alm T, Edqvist PH, Berling H, Tegel H, Mulder J, Rockberg J, Nilsson P, Schwenk JM, Hamsten M, von Feilitzen K, Forsberg M, Persson L, Johansson F, Zwahlen M, von Heijne G, Nielsen J, Pontén F. Proteomics. Tissue-based map of the human proteome. Science. 2015 Jan 23;347(6220):1260419. doi: 10.1126/science.1260419. PMID: 25613900.

Volkova M, Russell R 3rd. Anthracycline cardiotoxicity: prevalence, pathogenesis and treatment. Curr Cardiol Rev. 2011 Nov;7(4):214–20. doi: 10.2174/157340311799960645. PMID: 22758622; PMCID: PMC3322439.

von Gise A, Pu WT. Endocardial and epicardial epithelial to mesenchymal transitions in heart development and disease. Circ Res. 2012 Jun 8;110(12):1628–45. doi: 10.1161/CIRCRESAHA.111.259960. PMID: 22679138; PMCID: PMC3427736.

Zhang N, Yin Y, Xu SJ, Chen WS. 5-Fluorouracil: mechanisms of resistance and reversal strategies. Molecules. 2008 Aug 5;13(8):1551–69. doi: 10.3390/molecules13081551. PMID: 18794772; PMCID: PMC6244944.

